# Ketamine inhibition of SARS-CoV-2 replication in astrocytes is associated with COVID-19 disease severity in a variant-dependent manner

**DOI:** 10.1101/2025.03.29.646083

**Authors:** Borut Furlani, Maja Potokar, Miša Korva, Katarina Resman Rus, Patricija Pozvek, Tomaž M. Zorec, Julijan Vršnik, Samo Pirnat, Petra Tavčar Verdev, Tina Smolič, Anemari Horvat, Kristina Radinović, Nina Grasselli Kmet, Tatjana Avšič-Županc, Nina Vardjan, Matjaž Stenovec, Matjaž Jereb, Jernej Jorgačevski, Robert Zorec

## Abstract

Severe coronavirus infections, including SARS-CoV-2, can cause neurological symptoms, but the underlying neurotropic mechanisms are unclear. Experiments with SARS-CoV-2 variants B.1.258.17, B.1.1.7, and BA.5.3.2 (termed wild-type, alpha and omicron, respectively) revealed that human astrocytes, not neurons, support viral proliferation. During the COVID-19 pandemic, new virus variants exhibited milder disease progression. A retrospective study of patients with COVID-19 infected by wild-type or alpha variants was conducted to test whether ketamine, an anaesthetic that inhibits endocytosis, affects COVID-19. At admission, patients infected with the wild-type showed greater disease severity than alpha variant patients, but the disease course was similar. This may be due to distinct ketamine-mediated SARS-CoV-2 variant-dependent effects, revealing stronger ketamine inhibition of the wild-type variant than the alpha variant mediating astrocyte responses involving the expression of ACE2, a viral cell entry site, viral proteins RNA-dependent RNA polymerase and envelope protein-E in infected cells. Overexpression of SARS-CoV-2 protein 3a attenuated astroglial lysosomal traffic, and 3a and nsp6 differentially modulated lipid droplet accumulation and initiation of autophagy, where ketamine predominantly affected vesicle dynamics. In summary, human astrocytes, but not neurons, contribute to SARS-CoV-2 neurotropism, highlighting the potential benefits of ketamine treatment in coronavirus infections.

---

Severe acute respiratory syndrome coronavirus 2 (SARS-CoV-2) caused the Coronavirus disease 19 (COVID-19) pandemic. Neurological manifestations range from olfactory alterations to nausea, dizziness, and headaches, as well as encephalopathy and stroke^1^. The underlying mechanisms causing these signs and symptoms of the central nervous system (CNS) are unclear and may include hypoxia, immune-mediated damage, coagulation problems and viral invasion into the CNS^2^, which was noted to be very low and did not correlate with the severity of histopathological alterations^3^. Other post-mortem studies demonstrated the presence of SARS-CoV-2 viral RNA and proteins in various regions of the brain, indicating that SARS-CoV-2 enters the CNS, possibly via the olfactory epithelium^4^. In patients who died with COVID-19, the presence of SARS-CoV-2 was detected in ∼50% of brains examined, but this was not associated with the severity of disease^5^. A study examining a cohort of 26 individuals who died of COVID-19, using histopathological signs of brain damage as a guide for possible SARS-CoV-2 brain infection, revealed that among the five individuals who exhibited those signs, all of them had genetic material of the virus in the brain, exhibiting foci of SARS-CoV-2 infection and replication, particularly in astrocytes^6^.

Astrocytes have been considered suitable cells for viral replication. These homeostasis-providing cells^7^ exhibits aerobic glycolysis, a form of metabolism characteristic of proliferating and morphologically dynamic cells^8^. Consistent with this, ∼85% of patients who died of COVID-19 exhibited variable degrees of reactive astrogliosis^5^, a response to any pathologic insult^9,10^. We hypothesized that human astrocytes, but not neurons, can mediate SARS-CoV-2 infection. We exposed human astrocytes and neurons to SARS-CoV-2 variants and examined their capacity to support proliferation and further infection of neural cells. The results revealed that astrocytes, but not human neurons, have a much higher capacity to mediate SARS-CoV-2 infection with B.1.258.17 (wild-type), B.1.1.7 (alpha), and BA.5.3.2 (omicron) variants. The latter evolved more recently.

Ketamine, a dissociative anaesthetic^11^, has gained interest in mechanically ventilated patients with COVID-19, primarily indicated for its vasopressor and propofol-sparing effects^12^. Its use was shown to be feasible and safe as an adjunct analgo-sedative agent with no negative impact on patient outcome^13^. Ketamine is an anti-inflammatory molecule that could potentially contain the cytokine “storm” in severe COVID-19^14^. Molecular dynamics studies indicated ketamine to be a putative inhibitor of SARS-CoV-2 replication through the interaction of angiotensin-converting-enzyme 2 (ACE2), an enzyme mediating virus entry into cells^15^.

During the COVID-19 pandemic, each new emerging variant of SARS-CoV-2 has typically appeared with higher transmissibility and potential for immune escape among the population immunized against the older variants, but also with a milder disease severity^16,17^. To learn whether ketamine may be beneficial for the treatment of patients with COVID-19 hospitalized in the intensive care unit (ICU), a retrospective clinical study was conducted, where patients were sedated with ketamine, and the severity of disease at admission was determined by the APACHE (Acute Physiology and Chronic Health Evaluation) score^18^. At admission, patients infected with the wild-type variant had a more severe clinical presentation than those infected with the alpha variant, however, the course of COVID-19 disease appeared similar with both variants. Among many possibilities, this may be because ketamine inhibits human astrocyte infection with the wild-type more strongly than the alpha variant, which was investigated at the cellular level.

Astrocytes possess numerous mechanisms for viral entry, including the SARS-CoV-2 obligate receptor ACE2, linked to endocytosis^19,20^. We studied the sensitivity of this pathway to ketamine, which inhibits endocytosis^21^, likely through astrocyte-specific mechanisms^22^, by using human astrocytes infected with wild-type and alpha SARS-CoV-2 variants and quantitative microscopies, immunocytochemistry and molecular tests. We asked whether ketamine affects the SARS-CoV-2 infection rate by monitoring the percentage of infected cells, as well as the expression and cellular distribution of ACE2 receptor and two viral proteins (RNA-dependent RNA polymerase [RDRP] and envelope E protein [ENV]). Infection of astrocytes with either variant of SARS-CoV-2 upregulated the expression of ACE2, which was attenuated by ketamine. Furthermore, the subcellular densities of RDRP and ENV were lower in ketamine-treated SARS-CoV-2-infected astrocytes compared with the non-treated controls. Overexpression of SARS-CoV-2 proteins 3a and non-structural protein 6 (nsp6) revealed that 3a proteins attenuate lysosomal traffic and that both proteins modulate lipid droplet accumulation differently; although not affected by 3a, nsp6 stimulated lipid droplet accumulation. The 3a protein stimulated autophagy initiation, whereas nsp6 attenuated it. Both 3a and nsp6 decrease cell size, but ketamine predominantly inhibited vesicle dynamics. In summary, our study on human astrocytes highlights the potential benefits of ketamine treatment, likely acting through astrocytes, in patients with COVID-19.

## Results

### SARS-CoV-2 infectivity of human astrocytes exceeds that of human neurons

Whether the neurological signs and symptoms in patients with COVID-19 include entry of the SARS-CoV-2 into the CNS was addressed by several studies, which revealed inconsistent results^2–6^. To study the susceptibility of neural cells to SARS-CoV-2 infection directly, we infected human astrocytes and neurons with the B.1.258.17 (wild-type), B.1.1.7 (alpha) and BA.5.3.2 (omicron) SARS-CoV-2 variants at a multiplicity of infection of 0.1. Fig. 1 shows that astrocytes are more susceptible to infection with the SARS-CoV-2 variants than neurons; astrocytes were strongly immunolabelled with antibodies against RDRP (green), an enzyme mediating the proliferation of the virus (Fig. 1A_i_, _ii_). The infection of human astrocytes and neurons with the SARS-CoV-2 variants resulted in increased replication rates, quantified by the amount of viral RNA released in the supernatant using real-time reverse transcription polymerase chain reaction (RT-PCR). In cultured astrocytes, the replication rates of the SARS-CoV-2 variants were multiple orders of magnitude higher than in neurons (Fig. 1B_i__-iii_). We studied the tissue culture infectious dose 50 (TCID50), a parameter reporting the infectivity of the released infectious virus (i.e. 50% tissue culture infectious dose, determined in an endpoint dilution assay to measure infectious viral titre). In human astrocytes, the TCID50 of the SARS-CoV-2 variants increased with post-infection time, indicating efficient and productive replication of all the SARS-CoV-2 variants (Fig. 1 C_i-iii_, filled symbols). In contrast, in human neurons, the production of infectious viruses appeared low (Fig. 1C_i__-iii_, open symbols). These results confirm that human astrocytes are not only infected by SARS-CoV-2 and replicate the virus but that the virus released by astrocytes is also infective; the production of neuron-derived virus was negligible (Fig. 1B, C).

**Fig. 1.**
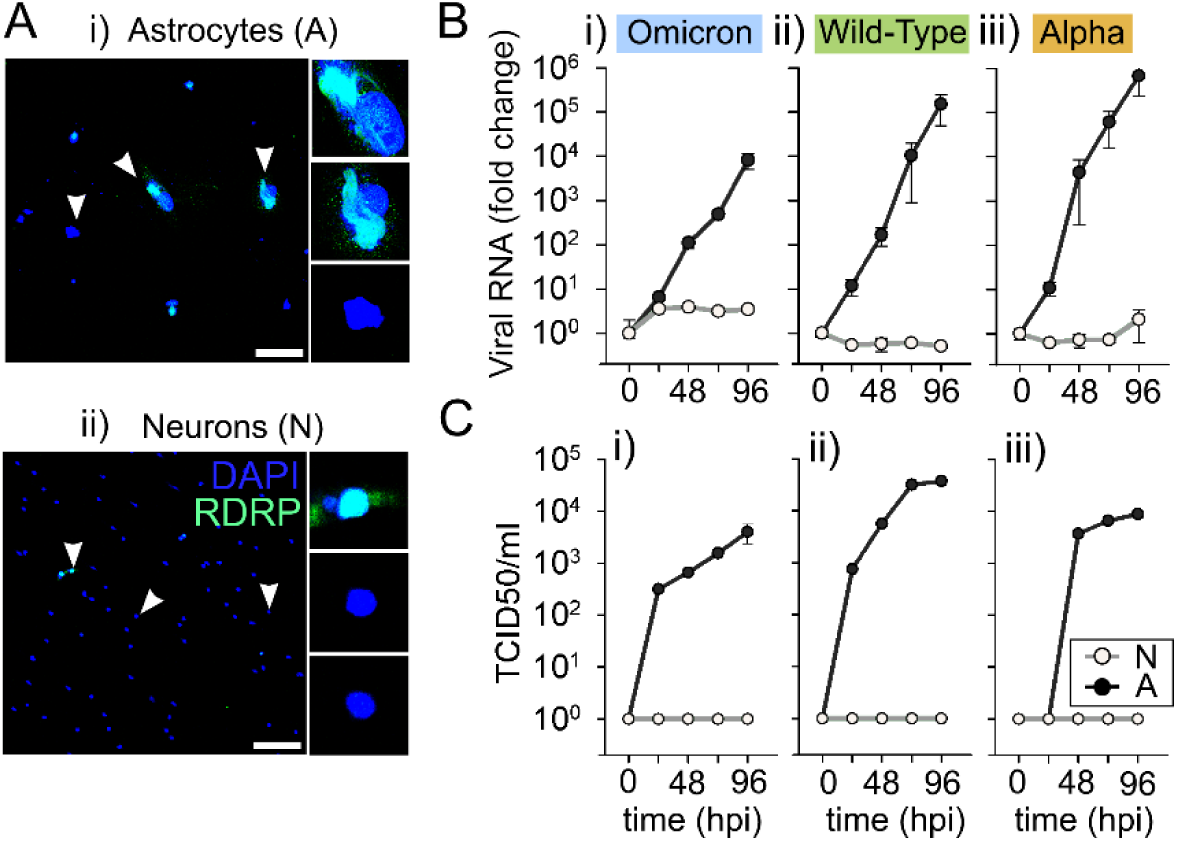
Human astrocytes are more susceptible to infection with SARS-CoV-2 than human neurons. (**A**) Human astrocytes (i) and human neurons (ii) infected with SARS-CoV-2 (omicron variant, BA.5.3.2) at multiplicity of infection of 0.1. Infected cells were immunolabelled with antibodies against RNA-dependent RNA polymerase (RDRP, green) at 48 h post-infection (hpi) and counterstained with the nuclear fluorescent dye DAPI (blue). Arrowheads point to the cells displayed in the enlarged insets. Notice the increased fluorescence intensity of infected (RDRP-positive) human astrocytes (i) compared with neurons (ii). (**B**) Replication rates of SARS-CoV-2 variants (i) omicron, (ii) wild-type (B.1.258.17) and (iii) alpha (B.1.1.7) in cultured human astrocytes (A, black symbols) and human neurons (N, open symbols), determined by quantifying the amount of viral RNA by real-time RT-PCR (see Methods) in the cell culture supernatant (viral RNA copies/ml) and (**C**) tissue culture infectious dose 50 (TCID50/ml, see Methods). In human astrocytes, the concentration of the RNA levels in all SARS-CoV-2 variants increased with hpi, indicating efficient and productive virus replication (black, B_i_, B_ii_ and B_iii_). In contrast, replication rates of all three SARS-CoV-2 variants were several orders of magnitude lower in neurons (open symbols; B_i_, B_ii_ and B_iii_). The TCID50 results shown in (**C**) confirm the infectivity of astrocyte-released virus (black; C_i_, C_ii_, C_iii_), whereas the production of neuron-derived infectious virus was absent (open symbols; C_i_, C_ii_, C_iii_). Data in (**B**) and (**C**) are presented on a logarithmic scale as means±standard error. Scale bar, 100 µm.

### At admission to the hospital, disease severity was higher in the group infected with wild-type, but the course of COVID-19 disease was similar in both groups

During the progress of the COVID-19 pandemic, every new variant of SARS-CoV-2 has typically appeared with milder severity. Human astrocytes are more prone to infection than neurons by various SARS-CoV-2 variants (Fig. 1), therefore, astrocytes are the likely mediators of neurological impairments in COVID-19. Ketamine, a dissociative anaesthetic^11^, is often used for sedation of critically ill patients. It was shown previously that ketamine treatment inhibits endocytosis by increasing cholesterol specifically in astrocytes^21–23^, and thiscould protect cells from infection by inhibiting virus entry. We asked whether ketamine sedation affects the course of COVID-19. This was studied in 96 patients infected with wild-type and alpha variants. Ketamine was applied as a bolus at the time of endotracheal intubation only (1–2 mg/kg of body weight) or continuously during the period of mechanical ventilation (up to 200 mg/h). The severity of the disease, rated using the APACHE score^18^, was determined on admission of patients. It was higher in the group infected with wild-type versus the alpha variant (Fig. 2A). However, the mortality rate was similar for both groups (Fig. 2B). Therefore, data from both ketamine-treated groups were pooled, and we next examined whether there were differences in the course of COVID-19. The median survival and hospitalization length of stay were similar for patients with wild-type versus alpha variants (Fig. 2C, D). In summary, at admission, patients infected with the wild-type variant exhibited higher disease severity, but the course of COVID-19 was similar in the two groups, indicating that ketamine may affect astrocytes infected with the wild-type more strongly.

**Fig. 2.**
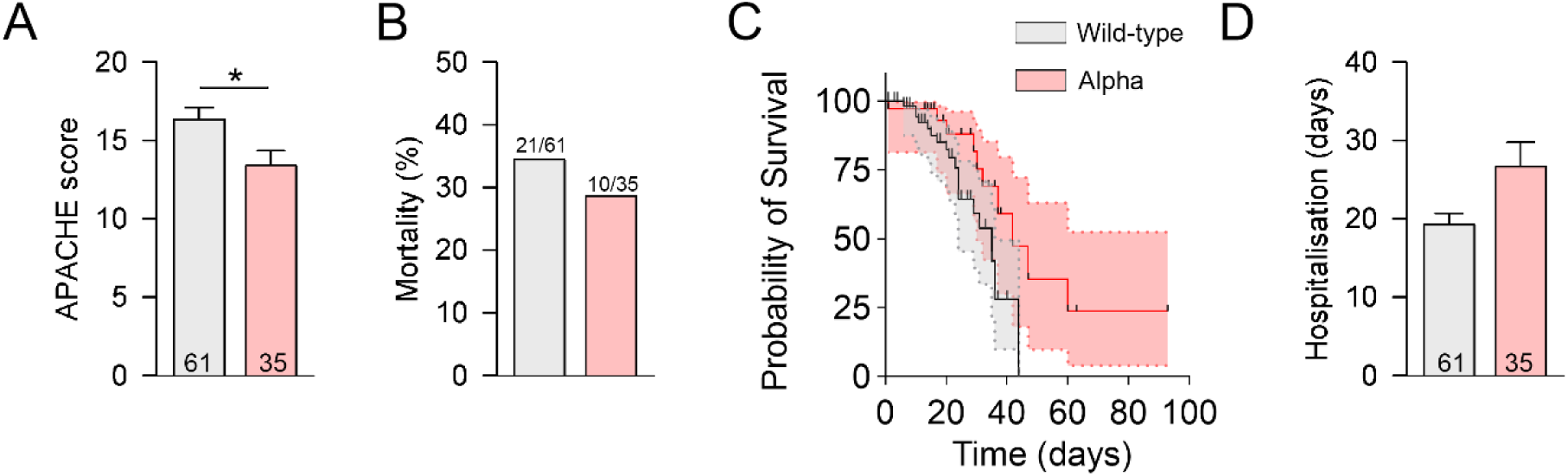
At admission to the intensive care unit (ICU), patients infected with the wild-type (B.1.258.17) variant of SARS-CoV-2 had a more severe clinical presentation than patients infected with the alpha variant (B.1.1.7), yet the course of the disease appeared similar. (A) Patients infected with the wild-type variant had a more severe clinical presentation than patients infected with the alpha variant, as determined by the APACHE score (B_i_, *P*=0.02, *t* test). (B) Mortality rates (determined as % of patients who died of COVID-19 in the ICU) were similar in both groups of patients. (**C**) Kaplan–Meier plot of overall survival with a 95% confidence interval (grey and red) in all patients who were infected with the wild-type (black) and alpha (red) variants, respectively. Survival time was measured from admission to the ICU until the date of death. Black symbols denote drop-out patients. Median survival times of patients infected with the wild-type were lower than those of patients infected with the alpha variant (35 versus 42 days, respectively, *P*=0.05, log-rank test). (**D**) Effect of SARS-CoV-2 variants on the mean duration of hospitalization. Patients infected with the wild-type variant exhibited slightly longer mean hospitalization times, however, the difference was not statistically significant (*P*=0.09, Mann-Whitney *U* test). All patients received treatment with ketamine and were mechanically ventilated. SARS-CoV-2 variants were determined according to the Pangolin classification system (Pango v.4.3.1). APACHE, Acute Physiology and Chronic Health Evaluation.

### Ketamine attenuated human astrocyte infection with the wild-type SARS-CoV-2 variant

Virus entry into cells engages many mechanisms, including endocytosis. We hypothesized that ketamine may affect SARS-CoV-2 infection, because it was shown that ketamine inhibits endocytosis in astrocytes^21^. We imaged RDRP and DAPI (4′,6-diamidino-2-phenylindole) fluorescent signals in human astrocytes (control and infected) using confocal microscopy (Fig. 3A). To evaluate infection rates, RDRP-positive (RDRP^+^) cells were counted and their percentage (the number of RDRP^+^ cells relative to all cells) was determined. The replication of SARS-CoV-2 was observed because several RDRP^+^ human astrocytes were counted in each field of view (Fig. 3A_iii_). Ketamine treatment reduced the percentage of RDRP^+^ in astrocytes infected with wild-type at 48 h post-infection (hpi; Fig. 3C). In part, this reduction could be due to ketamine-mediated cell death of astrocytes, and hence this may contribute to a reduction in virus replication. However, no time-dependent change in cell viability was noticed in controls. There was a gradual time-dependent decline in cell viability in SARS-CoV-2-infected cells, similar for both SARS-CoV-2 variants (Fig. 3B). These results are consistent with the view that ketamine treatment attenuates the rate of replication of wild-type SARS-CoV-2 in infected human astrocytes. Furthermore, to evaluate the release (production) of the viral RNA, we determined the concentration of viral RNA in the supernatant using RT-PCR (at intervals of 24 hpi from 0 hpi to 96 hpi). Both SARS-CoV-2 variants replicated similarly in human astrocytes (Fig. 3D). Under ketamine treatment, a significant reduction in SARS-CoV-2 RNA concentration was observed at 96 hpi in wild-type-infected cells (Fig. 3D_i_), but not in alpha-infected astrocytes (Fig. 3D_ii_). These results indicate differential sensitivity of human astrocytes infected with different variants of SARS-CoV-2 to ketamine treatment.

**Fig. 3.**
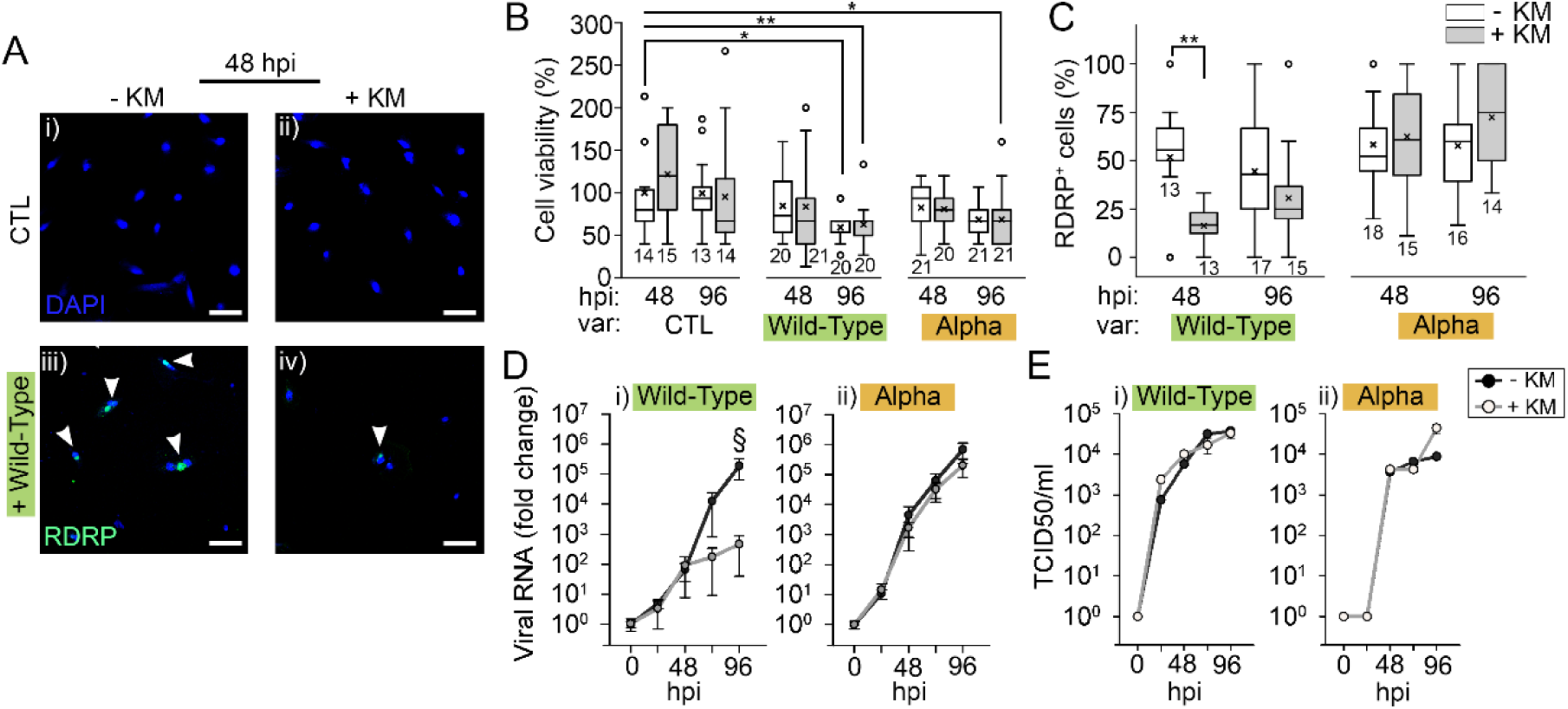
Ketamine treatment does not affect cell viability but reduces the susceptibility of human astrocytes to infection with the wild-type (B.1.258.17) variant of SARS-CoV-2. The number of cells, determined by counting DAPI-stained nuclei, after astrocyte infection with the wild-type variant in the absence (**A_i_**) or presence of ketamine (KM, **A_ii_**). SARS-CoV-2-infected cells were labelled by an antibody against viral RNA-dependent RNA polymerase (RDRP) in the absence (**A_iii_**) and presence of KM (**A_iv_**). Note that KM treatment reduced the percentage of RDRP-positive cells (**A_iii_** versus **A_iv_**). (**B**) Cell viability (%) calculated as the ratio between the average numbers of cells per field of view in infected (grey bars) and control (white bars) cells (CTL, non-infected, non-treated by ketamine). Treatment with ketamine did not affect astrocyte viability. (**C**) However, it significantly reduced the average percentage of RDRP-positive cells at 48 hpi in the case of the wild-type (B.1.258) variant (*P*=0.002, Mann-Whitney *U* test). (**D**) The concentration of viral RNA was measured in the supernatant at 24, 48, 72, and 96 hpi using a polymerase chain reaction (PCR). In human astrocytes infected with the wild-type variant, ketamine treatment reduced the viral RNA concentration at 96 hpi compared with non-treated controls (**D_i_**, \**p*<0.05, Mann-Whitney *U* test). In non-treated wild-type-infected cells, the viral RNA concentration increased exponentially (note log scale). In contrast, ketamine-treated astrocytes exhibited a slower increase in RNA concentration after 48 hpi (^#,§^*P*<0.05, ^##,§§^*p*<0.01, Wilcoxon rank sum test, compared with 0 hpi). The time course of SARS-CoV-2 RNA did not show differences between ketamine-treated and non-treated human astrocytes infected with the alpha variant (**D_ii_**). (**E**) The median tissue culture infectious dose (TCID50/ml) was similar in control and ketamine-treated human astrocytes infected with wild-type (**i**) or alpha (**ii**) variants of SARS-CoV-2. The numbers below the boxplots are the number of fields of view. CTL, control cells; hpi, hours post-infection; KM, ketamine; var, variant. Data are from two human donors. Scale bar, 100 µm (*^,#,§^*P*<0.05, **^,##,§§^*P*<0.01).

### Increased ACE2 receptor density in SARS-CoV-2-infected astrocytes is attenuated by ketamine

SARS-CoV-2 enters cells via the ACE2 receptor (Fig. 4A), although alternative entry pathways in astrocytes have been suggested^24^. ACE2 is expressed in astrocytic feet, components of the blood-brain barrier in human and rodent tissue^20^, with potential implications for neurological manifestations of brain ACE2 dysregulation during the acute or chronic phase of COVID-19 infection. We imaged subcellular ACE2 signals in z-stacks on a structural illumination microscopy (SIM; Fig. 4B_i_). The results support previous findings that human astrocytes express ACE2. Infection of human astrocytes with two SARS-CoV-2 variants significantly upregulated protein expression of ACE2 for both wild-type and alpha variants at 48 hpi, respectively (Fig. 4B_ii_, _iv_). However, the density of ACE2 was significantly lower after ketamine treatment in SARS-CoV-2-infected human astrocytes (Fig. 4B_iii_ and 4C, respectively). A similar reduction in ACE2 expression in astrocytes infected with the alpha variant was also observed at 96 hpi (Fig. 4C). No significant differences were noted among the controls.

**Fig. 4.**
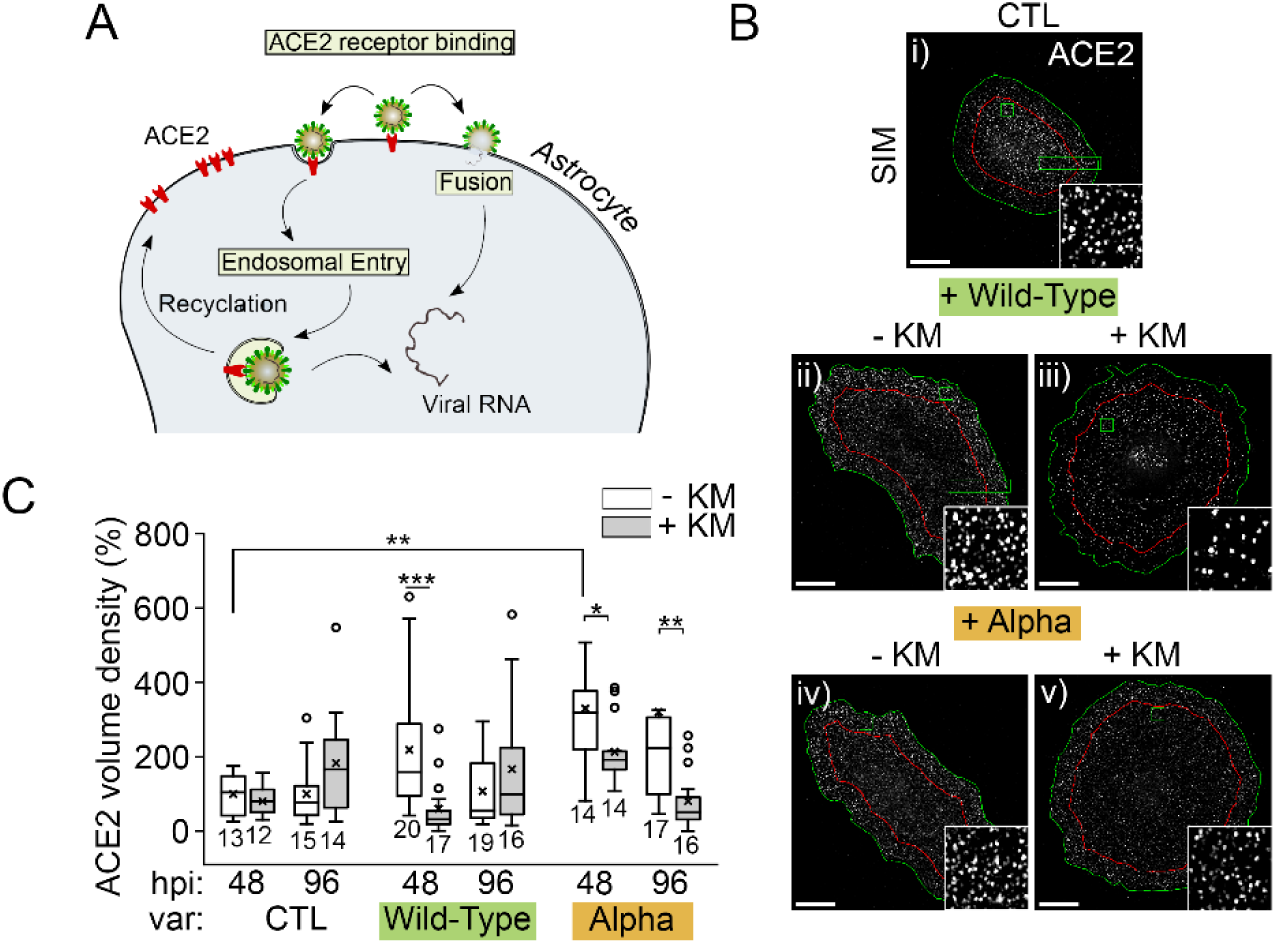
The density of ACE2 receptors is increased in human astrocytes infected with SARS-CoV-2, which is prevented by ketamine treatment. (A) The schematic illustrates two possible entry routes of SARS-CoV-2 into human astrocytes. Angiotensin-converting-enzyme 2 (ACE2) at the plasmalemma is considered the entry point into cells for SARS-CoV-2. After binding to ACE2, the virus is internalized either via the endosomal pathway or by direct fusion with the plasmalemma. (**B**) Human astrocytes were labelled with an antibody against ACE2 and observed with structured illumination microscopy. Infection with either SARS-CoV-2 variant resulted in an increase in the ACE2 particle density at 48 hpi compared with controls (**B_ii_** and **B_iv_**versus **B_i_**). However, ketamine treatment of infected human astrocytes resulted in a lower density of ACE2 compared with ketamine non-treated controls at 48 hpi (**B_iii_** versus **B_ii_**; and **B_v_** versus **B_iv_**), and in the case of the alpha variant, also at 96 hpi. (**C**) Observations were confirmed with quantitative analysis of the ACE2 signal density. Ketamine treatment reduced the ACE2 volume density in cells infected with the wild-type variant at 48 hpi (*P*<0.001, Mann-Whitney *U* test), and at 48 and 96 hpi in cells infected with the alpha variant (*P*=0.046 and *P*<0.001, Mann-Whitney *U* test, respectively). *n* is the number of cells. CTL, control cells; hpi, hours post-infection; KM, ketamine; SIM, structured illumination microscopy. The numbers below the boxplots are the number of cells analysed per experimental group. Data are from two human donors. Scale bar, 20 µm.

### Increased SARS-CoV-2 replication in human astrocytes leads to deformations in nuclear morphology, which are inhibited by ketamine

SARS-CoV-2 replication takes place in organelles that extend from the endoplasmic reticulum (ER), consisting of convoluted membranes and double-membrane vesicles (DMVs), associated with RDRP. SARS-CoV-2 infection resulted in morphologic remodelling of the endomembrane system and the nucleus (Fig. 5A). The SARS-CoV-2 replication organelles were visualized in human astrocytes by immunolabelled RDRP at 48 and 96 hpi. RDRP immunofluorescence was largely perinuclear, with scattered punctate structures throughout the cytoplasm (Fig. 5B, D), consistent with a previous report^25^. Ketamine treatment robustly decreased RDRP signal density in the case of the wild-type variant (Fig. 5B_ii_, C, D_iii_). In the case of the alpha variant, the effect was smaller (Fig. 5B_iv_, C). Furthermore, nuclei of SARS-CoV-2-infected human astrocytes underwent morphological deformations compared with controls, quantified by determining the shape factor (S.F., Fig. 5D)^26^. In control, non-infected astrocytes, ketamine did not affect the morphology of the nuclei. However, infection with either variant of SARS-CoV-2 resulted in a significant decrease in nuclear S.F. (Fig. 5D_i_ versus 5D_ii_, E). In astrocytes infected with wild-type, ketamine treatment recovered the S.F. to that measured in non-infected cells (Fig. 5D_iii_ versus 5D_i_, E). To determine the assembly and maturation of newly synthesized virions (Supplementary Fig. 1A), human astrocytes were immunolabelled for SARS-CoV-2 envelope (ENV) protein. ENV protein potentially serves as a marker of newly assembled/matured virions, however, it is also most likely present in endocytosed viral particles^27^. In human astrocytes infected with wild-type, ketamine treatment reduced the expression of ENV protein at 48 hpi (Supplementary Fig. 1Bii versus 1B_i_, C).

**Fig. 5.**
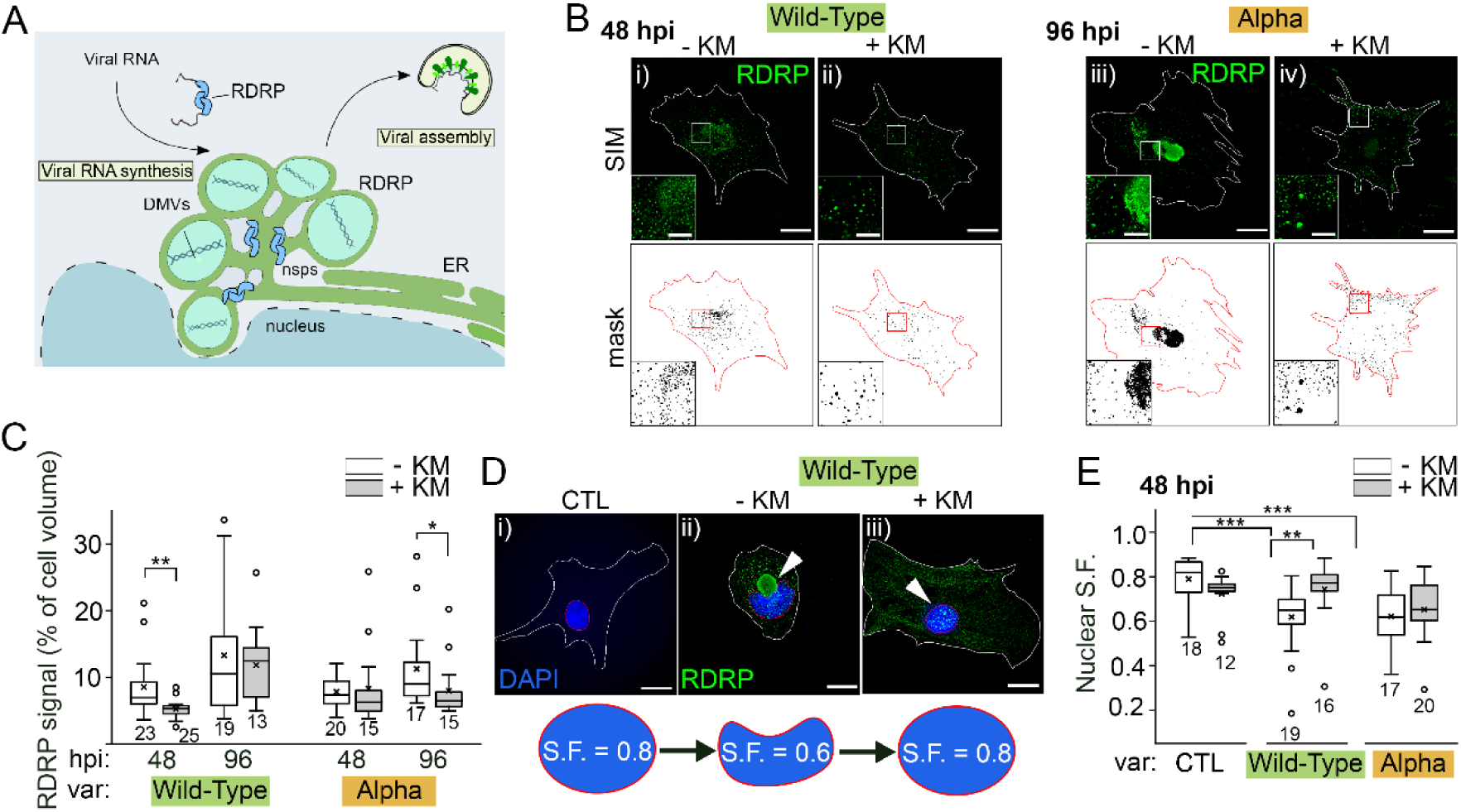
Ketamine treatment decreases the subcellular density of RNA-dependent RNA polymerase (RDRP) and restores nuclear deformations inflicted by SARS-CoV-2. (**A**) The schematic depicts the mechanism of SARS-CoV-2 replication in human astrocytes. (**B**) Fluorescent micrographs recorded by structured illumination microscopy show astrocytes immunolabelled against RDRP by anti-RDPR antibody (green). RDRP signal density was quantified on z-stack images of human astrocytes by determining the subcellular density, as a percentage relative to the RDRP signal that occupied the total cell volume. The average density of the RDRP signal was lower in ketamine-treated astrocytes infected with the wild-type-variant at 48 hpi (**B_ii_**), compared with non-treated controls (**B_i_**). Moreover, a similar reduction was observed in human astrocytes infected with the alpha variant at 96 hpi (**B_iii_** and **B_iv_**). (**C**) We quantified the average cell volume occupied by RDRP and confirmed that ketamine treatment reduced the density of RDRP in astrocytes infected with the wild-type variant at 48 hpi (*P*=0.003; Mann-Whitney *U* test), whereas a similar effect was observed in astrocytes infected with the alpha variant at 96 hpi (*P*=0.023; Mann-Whitney *U* test). The numbers below the boxplots are the number of cells. (**D**) Astrocyte nuclei were labelled with DAPI. In practically all cells, the RDRP signal formed a specific perinuclear structure, a place of extensive viral replication, which likely pressed against the cell nucleus resulting in nuclear deformation (**D_ii_**, arrowhead). Ketamine treatment partially inhibited the formation of the RDRP replication organelle (**D_iii_**, arrowhead). The circularity of the nuclei was characterized using a non-dimensional shape factor (S.F., see Methods), where 1 represents a circular nucleus and lower values indicate a non-circular morphology. (**E**) The analysis confirmed that infection with the wild-type and alpha variants reduced the S.F. (both *P*<0.001; Kruskal-Wallis test), whereas in astrocytes infected with the wild-type variant, ketamine treatment reversed the mean value of S.F. to that in the controls (*P*<0.001, Mann-Whiteny *U* test) (see schematic representation below the images). In contrast, ketamine treatment did not reverse the shape of the nuclei in astrocytes infected with the alpha variant. Data are from two human donors. CTL, control cells; DMV, double-membrane vesicles; ER, endoplasmic reticulum; hpi, hours post-infection; KM, ketamine; SIM, structured illumination microscopy; var, variant. Scale bar, 20 µm (insets, 5 µm).

### Expression of SARS-CoV-2 proteins in rat astrocytes reduces lysosome mobility, modulates lipid droplet accumulation and autophagy initiation, but ketamine predominantly attenuates vesicle dynamics

To further examine the action of ketamine in SARS-CoV-2-infected astrocytes, we transfected rat astrocytes to express the SARS-CoV-2 protein ORF3a (3a), which forms a large Ca^2+^-permeable, cation channel^28^, regulates autophagy^29^, and promotes lysosomal exocytosis and virion egress^30^. We also studied nsp6, a transmembrane protein that localizes to the ER membrane and the perinuclear space^31^. Among many processes, the nsp6 protein recruits lipid droplets (LDs) to replenish DMVs^28,29^ and impedes lysosomal acidification^31–34^.

### Protein 3a-mediated vesicle traffic is attenuated by ketamine

The SARS-CoV-2 protein 3a-EGFP (enhanced green fluorescent protein) predominantly segregates to acidified late endolysosomes (Supplementary Fig. 2)^30–32^. Lysosome mobility is reduced by elevating cytosolic calcium activity ([Ca^2+^]_i_)^35^. In this study, the mobility of vesicles labelled with LysoTracker Red DND-99 (LyTR; Thermo Fisher Scientific, Waltham, MA, USA) was measured, where LyTR is a marker of endolysosomes with acidified lumen^35^. We analysed control EGFP-and 3a-EGFP-expressing astrocytes (Fig. 6A, B and Fig. 6C, D, respectively). All vesicle mobility parameters, the total track length (TL, the length a vesicle travelled within 15 s), maximal displacement (MD, the maximal distance between the two positions in the vesicle track), directionality index (DI, the ratio of maximal displacement/total track length) and speed were significantly reduced in 3a-EGFP-expressing astrocytes versus the controls (Fig. 6E–H). Our data indicate that expression and/or structural association of 3a-EGFP with membranes of acidified late endolysosomes attenuated their mobility, most likely due to the increased resting levels of cytosolic Ca^2+^ (r[Ca^2+^]_i_; Fig. 6I–L), consistent with a previous report^35^. Moreover, increasing [Ca^2+^]_i_ by the application of ATP^35^ reduced TL and MD (Fig. 6M–P). We examined whether 10 µM ketamine, administered to astrocytes for 24 h, affects the mobility of 3a-EGFP vesicles that apparently deliver and likely incorporate protein 3a into the plasmalemma, where protein 3a may exert its pro-apoptotic activity^36^. Ketamine treatment reduced MD by 20% (Fig. 6R) and DI by ∼18% (Fig. 6G) compared to non-treated controls (Fig. 6R, S). Moreover, ketamine treatment impaired the incorporation of 3a-EGFP into the plasmalemma as revealed by reduced 3a-EGFP co-localization with the MemBrite 568/580 plasmalemma marker compared with non-treated controls (Supplementary Fig. 3).

**Fig. 6.**
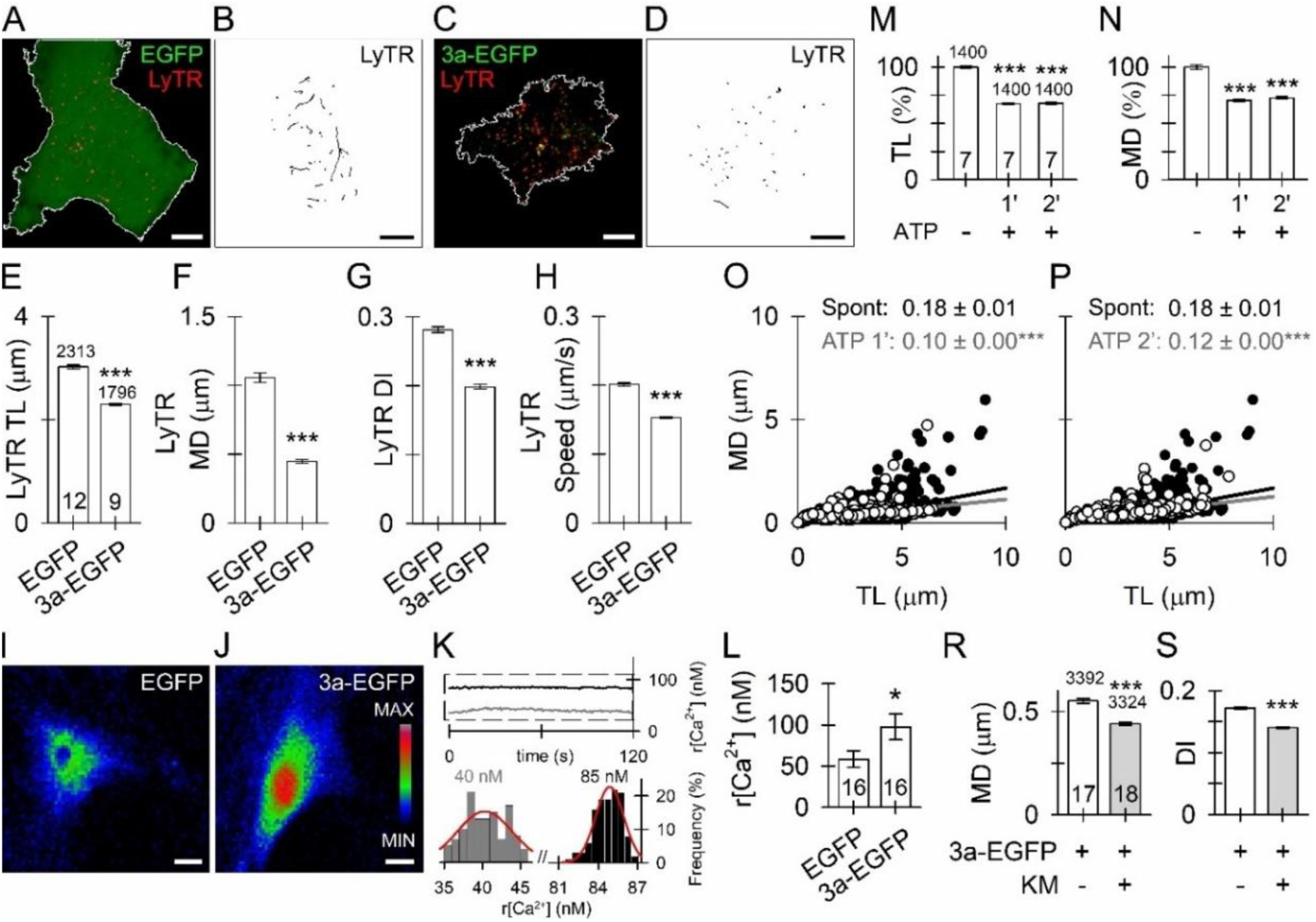
Expression of 3a-EGFP attenuates spontaneous mobility of LyTR-laden endolysosomal vesicles in rat cortical astrocytes, whereas ketamine further attenuates the mobility of 3a vesicles. (**A, C**) Double fluorescent confocal micrographs of LyTR-laden vesicles visible as red fluorescent puncta in the cytosol of EGFP-(**A**) or 3a-EGFP-expressing astrocytes (**C**); scale bars, 10 µm. (**B, D**) Reconstructed tracks of LyTR-laden vesicles (*n*=50) in a 15-s epoch. In 3a-EGFP-expressing astrocytes, LyTR-laden vesicles displayed limited mobility indicated by the contorted vesicle tracks (**D**), whereas in EGFP-expressing astrocytes, vesicle tracks were more elongated (**B**). (**E– H**) Plots displaying the track length (TL; **E**), maximal displacement (MD, **F**), directionality index (DI; **G**) and speed (DI; **H**) of LyTR-laden vesicles in astrocytes expressing EGFP (2313 vesicles; 12 cells) and 3a-EGFP (1796 vesicles; 9 cells). Note substantially attenuated mobility of LyTR-laden vesicles in 3a-EGFP-expressing astrocytes; \*\*\**P*<0.001 versus vesicle mobility in EGFP-expressing astrocytes (Mann-Whitney *U* test). (**I, J**) Epifluorescence images of transfected astrocytes expressing either EGFP (**I**) or 3a-EGFP (**J**) and loaded with the Ca^2+^ indicator Fura-2. The resting [Ca^2+^]_i_ is shown in a pseudo colour scale (right; minimum F340, black; maximum F340, red). Scale bars, 20 µm. (**K**) Examples of time-dependent change in [Ca^2+^]_i_ (nM) measured in resting astrocytes (black; r denotes resting intracellular Ca^2+^ concentration (r[Ca^2+^]_i_). The relative frequency distribution (%) of r[Ca^2+^]_i_ obtained during the 120-s resting period (indicated by a dashed box). These data were fitted with the Gaussian function, *f*=*a*×exp(−0.5×[(*x*−*x*_0_)/*b*]^2^), where *a*=15.29±2.27, *b*=3.59±0.82, *x*_0_=40.32±0.64 (grey frequency plot) in EGFP-expressing astrocytes and a=30.47±2.28, *b*=1.09±0.11, *x*_0_=84.86±0.10 (black frequency plot) in 3a-EFGP-expressing astrocytes. The parameter *x*_0_ is used to represent r[Ca^2+^]_i_ (shown in nM; r[Ca^2+^]_i_ values were calculated from the calibration curves). Note the higher baseline r[Ca^2+^]_i_ in 3a-EGFP-expressing astrocytes. (**L**) Average r[Ca^2+^]_i_ in EGFP-and 3a-EGFP-expressing astrocytes (data obtained from 16 cells from each culture). \**P*<0.05 (Mann-Whitney *U* test). (**M–P**) ATP stimulation, which increases astrocyte cytosolic calcium and suppresses the mobility of 3a-EGFP vesicles. (**M, N**) Mobility of 3a-EGFP vesicles (**M**, TL; **N**, MD; mean±SEM) before and during the first (1′) and the second (2′) minute after stimulation with 100 μM ATP. The numbers at the top and the bottom of the bars are the number of vesicles and cells analysed, respectively. \*\*\**P*<0.001 versus resting (non-stimulated) mobility (Mann-Whitney *U* test). (**O, P**) The plots display the relationship between MD and TL before (**O**, **P**, black circles), during the first (**O**, white circles), and the second (**P**, white circles) minute after ATP stimulation. A linear function (black and grey lines) of the form [MD=MD_0_+*a*×TL] was fitted to the data; the slopes (*a*±SEM) correspond to the directionality index (DI). This parameter (shown above the graphs) differed significantly (*P*<0.001; ANCOVA) within the first and the second minute after ATP stimulation. (**R, S**) Ketamine (KM) treatment suppresses the directional mobility of 3a-EGFP vesicles in astrocytes. Mobility of 3a-positive vesicles (**R**, MD; **S**, DI; mean±SEM) in non-treated (−) and ketamine-treated (+, KM, 10 µM) astrocytes. Significant reduction in vesicle MD and DI indicates hindered directionality of 3a-EGFP vesicles in cells. The numbers at the top and the bottom of the bars are the number of vesicles and cells analysed, respectively. \*\*\**P*<0.001 versus non-treated controls (Mann-Whitney *U* test).

#### nsp6, but not 3a-EGFP, triggered accumulation of lipid droplets in rat cortical astrocytes, which were associated with cell size reduction but not affected by ketamine

Unlike astrocytes expressing 3a-EGFP, nsp6-expressing astrocytes revealed an increase in the content of LipidTOX dye-labelled-LD, in particular an almost 2-fold increase in lipid droplet (LD) diameter (Fig. 7 [24 h] and Supplementary Fig. 4 [48 h]), suggesting a stress response and alterations in LD metabolism^37^. The nsp6-induced LD accumulation was prevented in the presence of selective inhibitors of diacylglycerol acyltransferase enzymes 1 and 2 (DGAT1 and DGAT2; Fig. 7G), responsible for triacylglycerol synthesis in ER ^38,39^, consistent with nsp6 recruiting LD for DMV formation^32^. nsp6-mediated LD accumulation in astrocytes was unaffected by ketamine treatment (Fig. 7I). In contrast, in 3a-EGFP-expressing astrocytes, there was a trend towards an increase in LD content, which was not significant compared with controls (Fig. 7D), but treatment with ketamine increased the diameter of LDs in 3a-EGFP-expressing astrocytes (Fig. 7D). Furthermore, we observed a decrease in cell area and cell perimeter in both nsp6-and 3a-EGFP-expressing astrocytes compared with controls (Fig. 7E, J). This suggests that both viral proteins might be involved in the recruitment of lipids from the plasmalemma for SARS-CoV-2-associated cellular processes, most likely for DMV formation^32^. The latter was not affected by ketamine treatment (Fig. 7). LDs were associated with both nsp6-and 3a-EGFP-labelled structures (Supplementary Fig. 5), consistent with previous studies^40^.

**Fig. 7.**
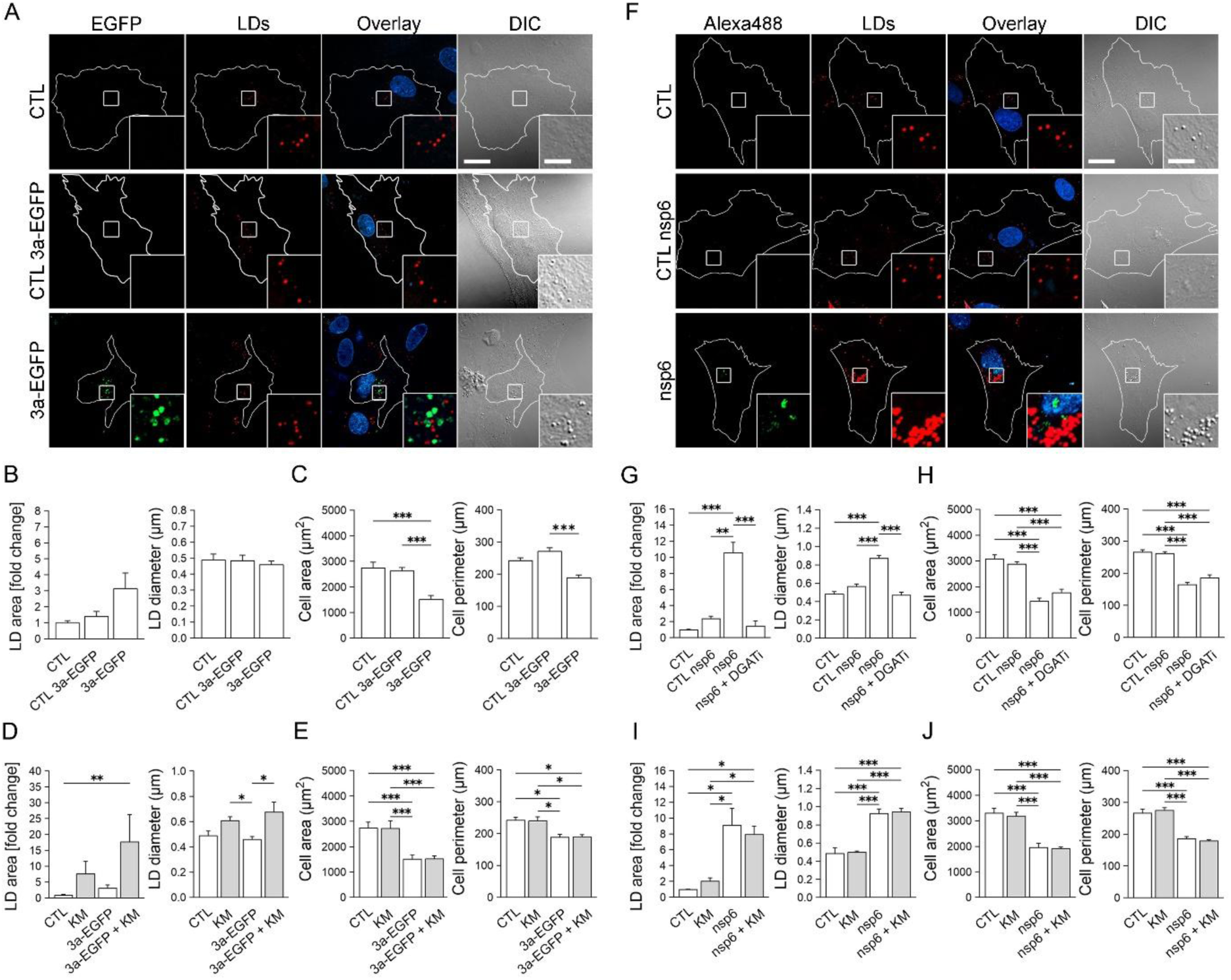
Lipid droplet dynamics in rat cortical astrocytes overexpressing SARS-CoV-2 3a and nsp6 proteins is independent of ketamine treatment. The EGFP-tagged 3a protein does not affect lipid droplet (LD) accumulation, but nsp6 enhances it; both mediate cell size reduction. (**A**) Representative fluorescence images of astrocytes in non-transfected and untreated cell cultures (CTL; *n*=7; 84 cells), in a cell culture transfected with pDNA encoding the viral EGFP-tagged 3a protein (3a-EGFP, green) either in a cell without apparent 3a expression (CTL 3a-EGFP; *n*=9; 116 cells) or in a cell expressing 3a (3a-EGFP; *n*=9; 151 cells) and labelled with fluorescent LD marker HCS LipidTOX red (red) and nuclear marker 4′,6-diamidino-2-phenylindole (DAPI, blue) 24 h after transfection. The overlay presents merged green, red and blue channels. DIC presents images taken by differential-interference contrast microscopy. The white line in the images shows the outline of an individual cell. Insets show white-boxed regions at higher magnification. Scale bars, 20 µm and 5 µm (inset). (**B**) Histograms of mean LipidTOX-stained cell area normalized to untreated controls (LD area [fold change]), mean diameter of LipidTOX-stained LDs (LD diameter), and (**C**) mean cell area and mean cell perimeter in CTL, CTL 3a-EGFP and 3a-EGFP astrocytes. Data are presented as means±SEM; *n*, number of independent experiments; \*\*\**P*<0.001 all pairwise (ANOVA, Holm-Sidak test [cell area] and ANOVA on ranks, Dunn’s test [cell perimeter]). (**D**) Histograms of mean LipidTOX-stained cell area normalized to control untreated samples (LD area [fold change]), mean diameter of LipidTOX-stained LDs (LD diameter), and (**E**) mean cell area and mean cell perimeter of cortical astrocytes in non-transfected and untreated cell cultures (CTL; *n*=7; 84 cells), in non-transfected cell cultures on 24-h exposure to 10 µM ketamine (KM; *n*=7; 84 cells), in cell cultures transfected with pDNA encoding the viral 3a protein (3a-EGFP; *n*=9; 151 cells) and in cell cultures transfected with pDNA encoding 3a on 24 h exposure to 10 µM KM (3a-EGFP+KM; *n*=8; 134 cells). Data are presented as means±SEM; *n*, number of independent experiments. \**P*<0.05, \*\**P*<0.01, \*\*\**P*<0.001 all pairwise (ANOVA, Holm-Sidak test [cell area] and ANOVA on ranks, Dunn’s test [LD area, LD diameter, cell perimeter]). (**F**) Representative fluorescence images of astrocytes in non-transfected and untreated cell cultures (CTL; *n*=7; 111 cells), in a cell culture transfected with pDNA encoding viral nsp6 protein, either in a cell without apparent nsp6 expression (CTL nsp6; *n*=15; 228 cells) or in a cell expressing nsp6 (nsp6; *n*=15; 244 cells), and nsp6-expressing astrocytes in a cell culture treated with DGAT1 and DGAT2 inhibitors (nsp6+DGATi; 10 µM; *n*=5; 56 cells). To trace nsp6 expression, LD formation and the presence of nuclei, cell cultures were labelled immunocytochemically with antibodies against nsp6 (Alexa Fluor 488, green), fluorescent LD marker HCS LipidTOX red (red) and nuclear marker DAPI (blue) 24 h on transfection. Overlays presents merged green, red and blue channels. DIC presents images taken by differential-interference contrast microscopy. The white line in the images shows the outline of an individual cell. Insets show white-boxed regions at higher magnification. Scale bars: 20 µm and 5 µm (inset). (**G**) Histograms of mean LipidTOX-stained cell area normalized to control untreated samples (LD area [fold change]), mean diameter of LipidTOX-stained LDs (LD diameter), and (**H**) mean cell area and mean cell perimeter in CTL, CTL nsp6, nsp6 and nsp6+DGATi astrocytes. Data are presented as means±SEM. *n*, number of independent experiments. \*\**P*<0.01, \*\*\**P*<0.001 all pairwise (ANOVA, Holm-Sidak test [LD diameter, cell area, cell perimeter] and ANOVA on ranks, Dunn’s test [LD area]). (**I**) Histograms of mean LipidTOX-stained cell area normalized to control untreated samples (LD area [fold change]), mean diameter of LipidTOX-stained LDs (LD diameter), and (**J**) mean cell area and mean cell perimeter of astrocytes in non-transfected cell cultures without treatment (CTL; *n*=3, 51 cells) and on 24-h treatment with 10 µM KM (*n*=4; 50 cells), and of nsp6-expressing astrocytes without treatment (nsp6; *n*=5; 62 cells) and on 24-h treatment with 10 µM KM (nsp6+KM; *n*=4; 52 cells). Data are presented as means±SEM; *n*, number of independent experiments; \**P*<0.05, \*\*\**P*<0.001 all pairwise (ANOVA, Holm-Sidak test).

#### nsp6 and 3a-EGFP attenuated autophagy in a ketamine-independent manner

To study whether ketamine affects autophagy, we immunolabelled pATG16L1, a marker of nascent autophagosomes Transfection of 3a-EGFP increased autophagy, but was unaffected by ketamine treatment (Fig. 8). In contrast, transfection of nsp6 reduced autophagy initiation, which was also unaffected by ketamine treatment (Fig. 8). Astrocytes that did not exhibit any visible expression of nsp6 also exhibited a decline in the pATG16L1 signal. These results revealed that protein 3a-and nsp6-associated autophagy processes appear insensitive to ketamine.

**Fig. 8.**
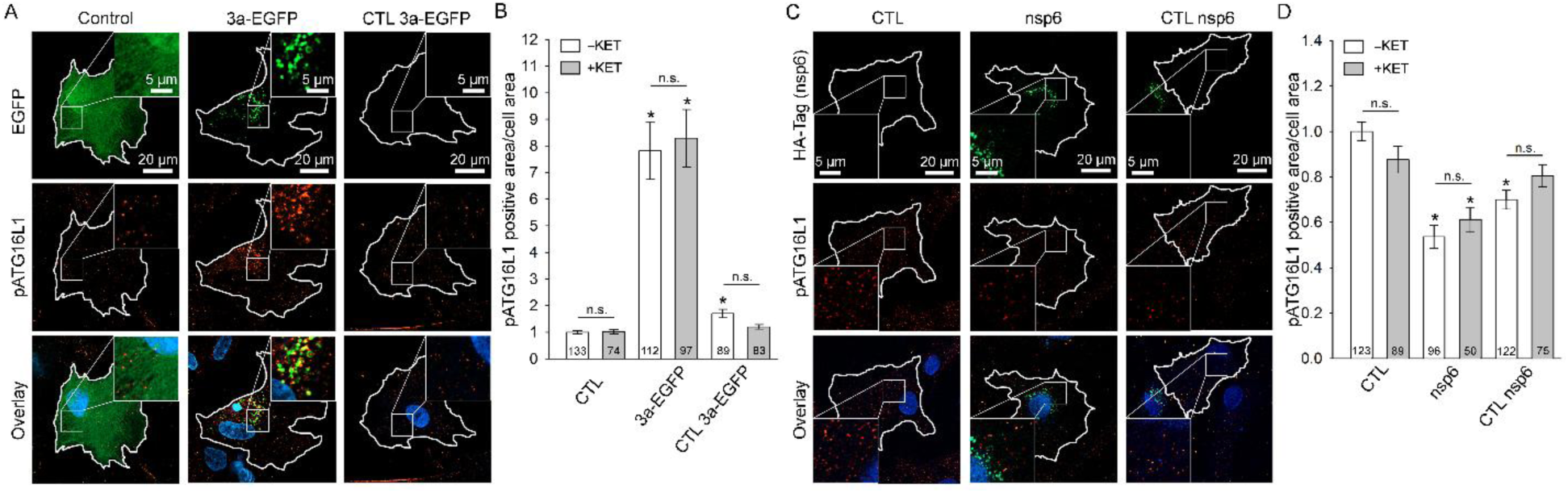
Autophagy initiation is modulated in cortical astrocytes overexpressing SARS-CoV-2 proteins 3a-EGFP and nsp6 independently of ketamine treatment. 3a-EGFP increases initiation of autophagy, but nsp6 strongly inhibits it. (**A**) Representative fluorescence micrographs of rat astrocytes expressing ORF3a (3a). Green fluorescence denotes either control cells expressing EGFP or cells expressing ORF3a-EGFP (3a-EGFP). Red fluorescence indicates immunolabelled pATG16L1 (a marker of nascent autophagosomes), and blue fluorescence denotes DAPI-stained cell nuclei. Cells of interest are outlined in white. Selected cell areas within the cell (white squares) are enlarged at the upper right corners of the respective micrographs. (**B**) Alterations in the autophagy initiation rate were determined by measuring the occupancy of the pATG16L1 signal (pATG16L1 positive area/cell area). Expression of 3a-induced initiation of autophagy and pATG16L1 signal occupancy increased in 3a-expressing cells compared with non-transfected cells in the same sample (3a-contrast) and EGFP-expressing control cells (Control). Treatment of astrocytes with ketamine did not affect the initiation of autophagy. \**P*<0.05 (ANOVA, Dunn’s test); n.s., not statistically significant. (**C**) Representative fluorescence micrographs of rat astrocytes expressing nsp6, immunolabelled with antibodies against nsp6 (HA-Tag; green fluorescence). Red fluorescence indicates immunolabelled pATG16L1 (a marker of nascent autophagosomes), and blue fluorescence denotes DAPI-stained cell nuclei. Cells of interest are outlined in white. Selected cell areas within the cell (white squares) are enlarged at the bottom left corners of the respective micrographs. (**D**) A histogram of alterations in the autophagy initiation rate was determined by measuring the occupancy of the pATG16L1 signal (pATG16L1 positive area/cell area). Expression of nsp6 inhibited the initiation of autophagy, as evident from a decrease in the occupancy of the pATG16L1 signal in nsp6-transfected cells (nsp6-transf) and non-transfected cells in the same sample (nsp6-nontransf), compared to non-transfected controls. Treatment of astrocytes with ketamine did not affect the initiation of autophagy. \**P*<0.05 (ANOVA, Dunn’s test); n.s., not statistically significant.

## Discussion

Here, we provide direct evidence that human astrocytes but not neurons mediate SARS-CoV-2 replication and further infection of neural cells, likely contributing to the neurological symptoms of COVID-19. Post-mortem studies that tested the presence of SARS-CoV-2 viral RNA and proteins in various brain regions, including the olfactory epithelium^4^, are in line with our results. Tissue damage was used as a guide for possible assessment of SARS-CoV-2 brain infection and replication, revealing that tissue damage is associated with astrocytes^6^. We used human astrocytes and neurons to test this hypothesis by directly infecting them with wild-type, alpha and omicron SARS-CoV-2 variants. The replication rates of the SARS-CoV-2 variants and the infectivity of the released virus in cultured human astrocytes increased with post-infection time and were orders of magnitude higher than in neurons. This indicated efficient and productive replication of the studied SARS-CoV-2 variants in astrocytes (Fig. 1), indicating that astrocytes appear as a key target for neurotropic viruses^19^, not only for SARS-CoV-2 (this study), but also for flaviviruses^41^, human immunodeficiency virus type 1 (HIV-1)^42^ and other virus types because astrocytes express several molecules and mechanisms mediating virus entry^19,43,44^ including endocytosis, found previously to be inhibited by ketamine^21^.

Disease severity was studied in ketamine-sedated patients infected by wild-type or alpha SARS-CoV-2 variants. Because the wild-type variant exhibited the highest TCID50 values in vitro (Fig. 1), we expected the clinical outcome in patients infected with these variants to be more severe than in those infected with the alpha variant. Disease severity at admission was higher in patients infected with the wild-type versus the alpha variant, monitored as the APACHE score. However, mortality rate, median survival and hospitalization length were similar in the two groups (Fig. 2), indicating that ketamine, a dissociative anaesthetic^11^, shown previously to specifically modulate astrocyte function and inhibit endocytosis^21,22^, may alleviate the severity of COVID-19, at least with the wild-type SARS-CoV-2 variant.

The similar course of COVID-19 disease in patients sedated with ketamine in our study likely reflects the distinct sensitivity to ketamine of astrocytes infected with the SARS-CoV-2 variants. This is supported by the molecular quantification of the viral RNA concentration measured in the supernatant, revealing that in human astrocytes infected with the wild-type SARS-CoV-2 variant, ketamine treatment reduced the viral RNA concentration at 96 hpi versus the non-treated controls (Fig. 3). Moreover, SARS-CoV-2 entry involves binding to ACE2 at the plasmalemma. After binding to ACE2, the virus is internalized either via the endosomal pathway or by direct fusion with the plasmalemma. In human astrocytes infected with the wild-type or alpha SARS-CoV-2 variant, the density of this receptor increased, but treatment with ketamine strongly inhibited SARS-CoV-2-induced increase in ACE2 receptor density (Fig. 4). To protect nascent viral genomic RNA from pattern recognition receptors that might trigger an innate interferon response, replication of the viral genome occurs in DMVs that originate from the ER to form replication organelles^45^. The average density of the RDRP signal was lower in ketamine-treated astrocytes infected with the wild-type (at 48 hpi) or the alpha SARS-CoV-2 variant (at 96 hpi), consistent with the inhibitory role of ketamine in astrocyte SARS-CoV-2 infection. We also observed that astrocyte nuclei, labelled with DAPI, were anatomically associated with the perinuclear RDRP structure, which was associated with nuclear deformation, as observed previously to be due to degradation of the DNA damage response checkpoint kinase 1 (CHK1), impaired 53BP1 recruitment and cellular senescence^46^. In our study, along with the partially inhibited formation of the RDRP replication organelle by ketamine, the circularity of the nuclei, characterized by a non-dimensional S.F., was also affected. Although the SARS-CoV-2 infection induced a decrease in S.F., ketamine treatment completely reversed the nuclear deformation in the case of the wild-type but not the alpha variant (Fig. 5). These results may underline the difference in COVID-19 disease severity observed in patients infected with the wild-type and alpha variant. Once the virions are formed, they exit the cell, usually via vesicle exocytosis. However, in the case of SARS-CoV-2, they are released from the infected cell through the lysosomal pathway^47^. To test whether ketamine affects this process, we expressed individual SARS-CoV-2 proteins in rat astrocytes to study lysosomal traffic, LD accumulation and autophagy initiation. In particular, we looked at protein 3a, a Ca^2+^ permeable channel^28^, shown to regulate autophagy^29^, and promote lysosomal exocytosis and virion egress^30^. We also studied nsp6, a transmembrane protein that localizes to the ER membrane and mediates i) the formation of replication organelles and DMVs that shield nascent viral RNA (vRNA), ii) antagonizes interferon type I (IFN-I) responses, iii) activates NLRP3 inflammasomes by impeding lysosomal acidification and iv) recruits LDs to replenish DMVs^31–34^.

The results revealed that expression and/or structural association of 3a-EGFP with membranes of acidified late endolysosomes attenuated their mobility. Previously, it was reported that increased [Ca^2+^]_i_ reduced vesicle mobility^35^. The increase in resting [Ca^2+^]_i_ observed here (Fig, is likely due to protein 3a channel-forming activity^28^ or to an application of ATP, releasing [Ca^2+^]_i_ from intracellular stores and reduced vesicle mobility parameters TL and MD (Fig. 6). Ketamine treatment reduced the mobility parameters by about one fifth relative to non-treated controls (Fig. 6). Although there are many mechanisms contributing to this phenomenon, ketamine action may be mediated via kinesin and by inhibition of endocytosis and cargo discharge from vesicles^21–23^. Ketamine treatment reduced protein 3a-EGFP co-localization with the MemBrite 568/580 plasmalemma marker (Supplementary Fig. 3), therefore, these results show that ketamine treatment hinders vesicle traffic and vesicle-mediated incorporation of protein 3a into the plasmalemma.

SARS-CoV-2 3a protein has been shown to trigger LD droplet accumulation in non-neural cells by blocking autophagic flux^48^, a process involved in the degradation of LDs^49,50^. However, although we observed that 3a is in contact with LDs in astrocytes (Supplementary Fig. 4), its overexpression did not significantly affect astroglial LD content (Fig. 7). Overexpression of the viral protein nsp6 in astrocytes significantly stimulated LD accumulation by increasing LD diameter, identifying nsp6 as a crucial regulator of LD metabolism for the first time. The accumulation of LDs in astrocytes triggered by nsp6 is dependent on the activity of key enzymes involved in LD biogenesis, DGAT1 and DGAT2, because inhibitors of these enzymes prevent LD accumulation (Fig. 7). This is consistent with observations in various non-neural cell lines infected by SARS-CoV-2, where inhibition of LD biogenesis not only reduces LD droplet accumulation but also decreases SARS-CoV-2 replication, indicating that LD are necessary for sustaining viral replication^40,49,51,52^.

The exact mechanisms behind nsp6-mediated LD accumulation in astrocytes remain to be fully elucidated. However, given that overexpression of nsp6 leads to inhibition of autophagy initiation (as shown in Fig. 8), this inhibition likely slows LD turnover, affecting their accumulation. In HeLa cells, nsp6 has been shown to zipper the ER membrane, facilitating lipid flow and restricting access of ER luminal proteins to DMVs. In addition, nsp6 was shown to act as a connector for LDs within the ER membrane, and these ER-LD connections were mediated through the LD-tethering complex DFCP1 (double FYVE domain-containing protein 1)–RAB18. Consistent with this, our results have shown that in astrocytes, nsp6 is in contact with LDs (Supplementary Fig. 4). This nsp6-mediated association of LDs with ER could provide fatty acids necessary for DMV biogenesis during viral replication in astrocytes, as observed in other non-neural cell types^31^. Supporting this, a decrease in astrocyte size was observed, suggesting that structural plasma membrane lipids are recruited to LDs, likely to fuel viral replication. nsp6-mediated LD accumulation is unaffected by ketamine treatment.

Autophagy initiation, involved in LD regulation^49^, was determined by immunolabelling of pATG16L1 in nascent autophagosomes (Fig. 8), revealing that overexpression of 3a-EGFP increased it, whereas nsp6 reduced it; both processes exhibited ketamine independence (Fig. 8). Astrocytes that did not show any visible expression of nsp6 exhibited a decline in the pATG16L1 signal, indicating that there may have been a trophic influence of nsp6-transfected cells onto the neighbor cells, mediating inhibition of autophagy initiation.

In conclusion, the results of this study provide direct evidence that human astrocytes but not neurons support SARS-CoV-2 proliferation and further infection of neural cells, most probably contributing to the neurological signs in patients with COVID-19. The retrospective clinical study of patients with COVID-19, sedated with ketamine and infected by wild-type or alpha variants revealed similar severity of disease between the two variants, which may reflect distinct ketamine sensitivity of astrocytes infected by the respective SARS-CoV-2 variants. In vitro studies confirmed that ketamine attenuates human astrocyte infection with the wild-type, but not the alpha SARS-CoV-2 variant. Furthermore, increased ACE2 receptor density in SARS-CoV-2-infected astrocytes, which facilitates further SARS-CoV-2 infection, is inhibited by ketamine, and ketamine treatment reduced SARS-CoV-2 replication and nuclear morphology deformations in human astrocytes. The mechanism of action of ketamine likely involves protein 3a-mediated processes, lysosomal traffic, which are part of LD and autophagy regulation. Ketamine inhibition of SARS-CoV-2 replication in astrocytes is therefore reflected in the disease course of COVID-19 in a variant-dependent manner, highlighting the benefits of ketamine treatment for patients with COVID-19.

## Online methods

Unless stated otherwise, all chemicals were obtained from Sigma-Aldrich (Merck KGaA), Darmstadt, Germany.

### Cell cultures and solutions

Cryopreserved primary human astrocytes were purchased from Innoprot (P10251, P10254). After thawing, cells were grown in cell culture flasks for 4 days and then plated on glass coverslips coated with poly-D-lysine (diameter 22 mm; Sigma-Aldrich, P6407) at a cell density of 1.5 ×10^4^ cells per coverslip (for immunocytochemistry experiments) or on 12-well culture plates (TPP, 92112) at a cell density of 3 ×10^4^ cells per well (RT-PCR experiments). Cell numbers were measured using a Scepter cell counter (Merck KGaA, Darmstadt, Germany). Cell cultures were maintained in standard astrocyte growth medium high-glucose Dulbecco’s modified Eagle’s medium (DMEM; Sigma-Aldrich, D5671) supplemented with 10% fetal bovine serum (FBS; Sigma-Aldrich, F7524), 1 mM sodium pyruvate (Sigma Aldrich, S8636), 2 mM L-glutamine (Sigma Aldrich, G3126), 5 U/ml penicillin and 5 µg/ml streptomycin (Sigma-Aldrich, P0781) at 37°C in a 5% CO_2_ atmosphere with 95% relative humidity. Cells were used in experiments approximately 24 h after plating on coverslips/culture plates. Primary human astrocytes were checked for cell culture purity by GFAP (Glial Fibrillary Acidic Protein) and ALDHL1 (Aldehyde Dehydrogenase 1 Family Member L1) immunostaining, revealing a purity of astrocytes >95%.

Primary cultures of rat cortical astrocytes were prepared from the cortex of neonatal Wistar rats (2–3 days old) as described previously^53^. Isolated cells were maintained in high-glucose DMEM with 4500 mg/l D-glucose), supplemented with 10% FBS, 1 mM sodium pyruvate, 2 mM l-glutamine, 5 U/ml penicillin, and 5 µg/ml streptomycin in an atmosphere of 5% CO_2_/humidified air (95%) at 37°C. Sub-confluent cultures were shaken at 225 rotations per minute overnight with a subsequent medium exchange; this was repeated three times. After enrichment, astrocytes were detached from the culture flask with 0.1% trypsin and 0.04% EDTA in Hank’s balanced salt solution, plated onto poly-L-lysine-coated 22-mm glass coverslips and bathed with the culture medium. Experiments were carried out 1–4 days after cell plating when cells reached 60–70% confluence to avoid confluence-induced generation of LDs^54^.

Extracellular solution (ECS) consisted of 130 mM NaCl, 5 mM KCl, 2 mM CaCl_2_, 1 mM MgCl_2_, 10 mM D-glucose, and 10 mM HEPES (4-(2-hydroxyethyl)-1-piperazineethanesulfonic acid), pH 7.2 with NaOH. Solution osmolarity (300 ± 15 mOsm) was measured with a freezing-point osmometer (Osmomat 030, Gonotec, Berlin, Germany). ATP was added to the extracellular solution as a bolus to reach a final concentration of 100 µM.

Animal handling and experimental procedures followed the International Guiding Principles for Biomedical Research Involving Animals developed by the Council for International Organizations of Medical Sciences, Animal Protection Act (Official Gazette of the Republic of Slovenia no. 38/13) and the Directive on Conditions for Issue of License for Animal Experiments for Scientific Research Purposes (Official Gazette of the Republic of Slovenia no. 40/85 and 22/87). The experimental protocol was approved by the Administration of the Republic of Slovenia for Food Safety, Veterinary and Plant Protection (Republic of Slovenia, Ministry of Agriculture, Forestry and Food, Dunajska cesta 22, 1000 Ljubljana), document no. U34401-47/2014/7 and U34401-48/2014/7, signed by Barbara Tomše, DVM.

### Astrocyte transfection and fluorescent labelling of astrocyte vesicles and plasmalemma

To visualize and examine the subcellular distribution of EGFP and protein 3a, we transfected cultured astrocytes with the plasmid encoding EGFP or wild-type SARS-CoV-2 protein 3a tagged with enhanced green fluorescent protein (3a-EGFP^28^). Astrocytes were transfected with pGBW-m4134157 plasmid (Addgene plasmid #152062) encoding for viral protein SARS-CoV-2 nsp6 protein with hemagglutinin (HA) tag, pSBL_DK186 plasmid encoding for insert SARS-CoV-2 3a protein fused to EGFP (Addgene plasmid #156422), or EGFP plasmid, kindly provided by Dr. Nicolas Vitale, CNRS, INCI, Strasbourg, France, using FuGENE transfection reagent (Promega, Madison, USA) according to the manufacturer’s instructions. Briefly, 3 µl of FuGENE 6 (Promega Corporation, Madison, WI, USA) was diluted in 100 µl of culture medium, mixed and incubated for 5 min at room temperature (RT). Next, DNA (0.8 µg/coverslip) was added, mixed and incubated for a further 15 min at RT. Astrocytes were washed and subsequently incubated in 900 µl of fresh culture medium to which 100 µl of the lipofection mixture was added. Transfected astrocytes were incubated for 24 h at 37°C in an atmosphere of 5% CO_2_/95% air. The medium was exchanged for fresh culture medium the next day, and transfected cells were observed microscopically after 24–48 h. In ketamine-treated astrocytes, ketamine was added to the growth medium to reach a final concentration of 10 µM for 24 h.

In a subset of experiments, acidic endosomes/lysosomes were visualized in EGFP– (controls) and 3a-EGFP–transfected cells by the addition of 200 nM of LysoTracker Red DND-99 (LyTR, Thermo Fisher Scientific) to the culture medium for 5 min at 37°C before the onset of experiments. The LyTR was excited by a 561-nm diode-pumped solid-state (DPSS) laser line and emission fluorescence was filtered with a bandpass filter 565–615 nm. In another subset of experiments, 3a-EGFP–transfected cells were supplied with 400 μl of ECS containing 4 μM FM4-64 (T3166, Thermo Fisher Scientific) to stain the plasma membrane and vesicles internalized by endocytosis^55^. FM4-64 was excited by a 488 nm argon laser line and emission fluorescence was filtered with a bandpass filter 630–755 nm.

### Calcium imaging and analysis

Fura-2 calcium measurements were performed as described previously^56^. In brief, 48 h after transfection with either pEGFP or p3a-EGFP, astrocyte-loaded coverslips were incubated for 30 min at RT in 1 ml of growth medium supplemented with 4 μM Fura-2 AM, a cell-permeant ratiometric fluorescent calcium indicator^57^. Next, cells were washed twice and incubated in fresh medium for an additional 30 min at RT. Cells were then washed twice, supplied with 400 μl of ECS and mounted in the recording chamber on an inverted fluorescence microscope (Axio Observer.A1; Zeiss, Jena, Germany). Cells were imaged using an EC Plan Neofluar 10×/NA 0.3 objective. To identify 3a-EGFP-or EGFP-expressing cells, we first recorded single images using a dye (excitation at 488 nm) and band-pass filtered the emission fluorescence at 515–565 nm. Fura-2 was excited by a monochromator Polychrome V (Till Photonics, Graefelfing, Germany) using filter set 21 HE (Zeiss), the emission fluorescence was band-pass filtered at 420–600 nm and captured by a CCD camera Sensicam QE (PCA AG, Kelheim, Germany). During the measurements, cells were sequentially excited at 340 nm and 380 nm (100-ms exposure with 10-ms time delay between the excitation wavelengths) at 1-s intervals for a total of 2 min. For analysis, we drew regions of interest (ROIs) delimiting cell soma and calculated the ratio of fluorescence intensities (F_340_/F_380_) recorded with Till Photonics Live Acquisition software corrected for background fluorescence. Ratio signals we calibrated as described previously^56^.

### Analysis of vesicle mobility

Astrocyte-loaded coverslips were mounted into the recording chamber and transferred to a confocal microscope (LSM 780; Zeiss) equipped with a plan-apochromatic oil-immersion objective 63×/NA 1.4. 3a-EGFP was excited using a 488-nm argon laser line and emission fluorescence was filtered with a bandpass filter (495–530 nm); LyTR was excited using a 561-nm DPSS laser line and emission fluorescence was filtered with a band-pass filter (565–615 nm). Time-lapse images were acquired at a rate of 2 Hz for 1 min. Vesicle mobility was analyzed in exported tiff files with the ParticleTR software (Celica Biomedical) as described previously ^58,59^. The mobility of 3a-EGFP and LyTR-laden vesicles was analyzed in exported tiff files with the ParticleTR software (Celica Biomedical, Ljubljana, Slovenia)^58^. Typically, 50 randomly selected 3a-EGFP or LyTR-laden vesicles were selected per transfected astrocyte, and their movement was tracked automatically as long as they remained in the focal plane while moving. Whenever the tracked vesicle moved into proximity to another vesicle, the automatic tracking was halted and the position of the selected vesicle was determined manually until the tracked vesicle separated enough from the neighbouring vesicle to resume the automated tracking. The following mobility parameters were determined for 15-s epochs: TL (total length of the travelled pathway), MD (maximal displacement; the farthest translocation of a vesicle), DI (directionality index; the ratio of MD/TL) and speed^58^.

### Determination of SARS-CoV-2 variants in samples

DeterminationSARS-CoV-2 variants in clinical samples was performed as previously described^60^. Briefly, sequence data were recorded on an Illumina NextSeq550 instrument (Illumina, San Diego, CA, USA); complete viral genome amplicon sequencing libraries were prepared using the commercially available COVIDSeq library preparation kit (Illumina). Sample-consensus complete SARS-CoV-2 genome sequences were called from maps of sequencing reads aligned to viral reference sequence Wuhan-Hu-1 (GenBank accession no. NC_045512.2), after careful quality and adapter trimming/filtering (QC) procedures. Briefly, the genome analysis pipeline used the Bbduk script (BBTools, http://sourceforge.net/projects/bbmap/) for QC procedures, Bwa Mem (http://arxiv.org/abs/1303.3997) for read mapping and Ivar was used for consensus calling (https://doi.org/10.1101/383513). The genome reconstruction pipeline was implemented as part of the Spike Screen pipeline, see the Github repository https://github.com/IMIMF-UNILJSI/scov2-spikeScreen.git for detailed parameter settings.

SARS-CoV-2 variant designations were assigned according to the Pangolin lineage classification system (https://doi.org/10.1038/s41564-020-0770-5) using the Pangolin command line tool (https://github.com/cov-lineages/pangolin.git, https://doi.org/10.1093/ve/veab064).

Experiments involving viruses were carried out in a biosafety level 3+ facility. The following characterized SARS-CoV-2 variants were used: wild-type (B.1.258.17; EVA-global portal access number: 005V-04394, Gisaid ID EPI_ISL_17777046), alpha (B.1.1.7; EVA-global portal access number: 005V-04053, Gisaid ID EPI_ISL_877453) and omicron (BA.5.3.2; EVA-global portal access number: 005V-04726, Gisaid ID EPI_ISL_17777060).

### Real-time RT-PCR

To quantify SARS-CoV-2 RNA with real-time RT-PCR, supernatants were collected at 0, 24, 48, 72, and 96 h. Total nucleic acid was extracted from 300 µl of cell culture supernatants with the OptiPure Viral Auto Plate, Proteinase K Reagent Kit (TANBead, Taoyuan City, Taiwan) on an automated nucleic acid isolation instrument Maelstrom 9600 TANBead. Viral RNA was quantified with LightMix™ SARS-CoV-2 E+N UBC (CE-IVD) using real-time RT-PCR on a QuantStudio 7Pro Real-Time PCR System (Thermo Fisher Scientific). Assays were performed in duplicate on cells from at least two human donors (purchased from Innoprot, see section on Cell cultures). Ketamine was added to the culture media in experiments studying the role of ketamine in SARS-CoV-2 replication. To account for donor-and experiment-specific variability, PCR data were normalized by dividing each timepoint by the donor-specific t_0_ average, and expressing results as fold changes relative to baseline. Normalized data were pooled for the subsequent analysis.

### TCID50

Infectious virus was determined by calculating TCID50. Briefly, Vero E6 cells (ATTC CRL-1586) were seeded at the concentration of 10^5^/well in 96-well plates (TPP, 92196), with the growth medium DMEM GlutaMAX (Thermo Fisher Scientific, 61965026) and 10% FBS (Euroclone, ECS0180L) and incubated for 24 h at 37° C in 5% CO_2_. Then, 50 µl of the medium was removed from the cells and supplemented with tenfold serial dilutions of suspension collected from the supernatant of virus-infected human astrocytes in 8 replicates. Plates were incubated at 37°C with 5% CO_2_ after 5 days. Plates were then inactivated with 100 µl of 4% formaldehyde solution for 30 min. Inactivated plates were visually inspected under the microscope for the presence of a cytopathic effect. TCID50 was calculated using the Spearman & Kärber algorithm^61^.

### Immunocytochemistry

Immunocytochemical labelling was performed as described previously (Potokar et al., 2014). Cells were briefly washed in phosphate-buffered saline (PBS; Sigma-Aldrich, P4417) and then fixed in 4% formaldehyde in PBS solution for 15 min at RT. Cells were then permeabilized with Triton X-100 (Merck Millipore, 1086031000) for 10 min at RT. After fixation and permeabilization cells were washed with PBS and incubated in 10% goat serum (Sigma-Aldrich G6767) and 3% bovine serum albumin (BSA; Sigma-Aldrich, A2153) solution in PBS for 1 h at 37°C to reduce non-specific binding of antibodies. Non-specific background staining was reduced by incubating the cells in a blocking buffer containing 3% BSA and 10% goat serum in PBS at 37 °C for 1 h. After the blocking step cells were incubated with primary antibodies against angiotensin-converting-enzyme 2 (Abcam, ab89111, dilution 1:50) and against two SARS-CoV-2 antigens: RNA-dependent-RNA-polymerase nsp12 (Genetex, GTX135467, dilution 1:100) or envelope N-term (ProSci, 3531, dilution 1:100) for 2 h at 37 °C or overnight at 4 °C. The cells were then rinsed in PBS and incubated in secondary antibodies against mouse or rabbit IgGs coupled to Alexa Fluor 546 (Invitrogen, Thermo Fisher Scientific, A110033; dilution 1:600) and Alexa Fluor 488 (Invitrogen, Thermo Fisher Scientific, A11008, dilution 1:600) for 45 min at 37°C (the mixture of secondary antibodies was applied simultaneously). After incubation, the cells were washed in PBS and mounted using SlowFade Gold antifade reagent with DAPI (Thermo Fisher Scientific, S36942) to diminish fluorescence bleaching and to stain cell nuclei, respectively. Control samples stained with secondary antibodies only were prepared in parallel to determine non-specific staining.

To examine the subcellular distribution of 3a-EGFP-positive structures in astrocytes, we quantified fluorescence co-localization between 3a-EGFP and immunocytochemically labeled proteins of endosomal compartments (i.e. early endosomes, autophagosomes, late endosomes/lysosomes). Transfected cells were washed (3 min) with PBS and fixed in formaldehyde (2% in PBS) for 10 min, permeabilized with 0.1% Triton X-100 for 10 min, and then washed four times with PBS at RT. The non-specific background staining was reduced by incubating cells in a blocking buffer with 10% (v/v) goat serum in PBS for 1 h at 37°C. Then cells were washed four times with PBS and incubated with primary antibodies diluted in 3% (w/v) BSA in PBS overnight at 4°C. The following primary antibodies were used: rabbit monoclonal anti-Rab7 (1:200; ab137029, Abcam), rabbit monoclonal anti-pATG16L1 (1:100; ab195242, Abcam), rabbit polyclonal anti-TPC1 (two-pore channel 1, clone 3526#6C, 1:500; a gift from Prof. Dr. Norbert Klugbauer, Albert-Ludwigs-University, Freiburg, Germany). The next day, the cells were washed in PBS (4 × 3 min) and stained with secondary anti-rabbit or anti-mouse antibodies conjugated to Alexa Fluor 546 (1:600; Thermo Fisher Scientific) at 37°C for 45 min. The cells were then washed in PBS (4 × 3 min) and mounted onto glass slides using SlowFade Gold antifade (Thermo Fisher Scientific) without DAPI.

In a subset of experiments, transfected control or ketamine-treated (10 µM for 24 h) astrocytes were stained with a fixable cell surface staining kit (MemBrite Fix 568/580; MBF568; 30095-T, Biotium, Fremont, CA, USA) according to the manufacturer’s instructions. Astrocyte-loaded coverslips were first incubated in the pre-staining solution (1×; 10 min at 37°C) and then in the staining solution (1×; 5 min at 37°C). Next, stained cells were washed in PBS (2× for 3 min), fixed (4% paraformaldehyde, 20 min, RT), washed again and mounted onto glass slides. Double-fluorescent cells were observed with a confocal microscope (LSM 780; Zeiss) with a plan-apochromatic oil-immersion objective 63×/NA 1.4. Z-stacked images were obtained with a 488-nm argon laser and 561-nm DPSS laser excitation, and the fluorescence emission was bandpass filtered at 495–530 nm and 565–615 nm, respectively. To quantify the co-localization of green-emitting (3a-EGFP) and red-emitting fluorophores (immunofluorescent Rab7, pATG16L1, TPC1 and MBF568), 8-bit TIFF files were exported and analysed with ColocAna (Celica Biomedical, Ljubljana, Slovenia), which enables automated high throughput co-localization analysis of fluorescent markers in a large number of images^62^. Briefly, the programme counted all green, red and co-localized (green and red) pixels in each image. The fluorescence co-localization (%) of red immunolabelled markers was determined for the green 3a-EGFP pixels. The threshold for the co-localized pixel count was set to 20% of the maximal fluorescence to minimize fluorescence overlap originating from closely positioned fluorescent structures, and structures above and below the focal plane that contribute some out-of-focus light scatter. In live cell labelling, fluorescence co-localization between red-emitting FM4-64 (630–755 nm), Dex546 and LyTR (565–615 nm), and green-emitting 3a-EGFP was quantified as stated earlier.

### Lipid droplet experiments

#### Astrocyte transfection and stimulation

Twenty-four hours after seeding, astrocytes were transfected with pGBW-m4134157 plasmid encoding for viral protein SARS-CoV-2 nsp6 protein with HA-Tag or pSBL_DK186 plasmid encoding for insert SARS-CoV-2 3a protein fused to EGFP using FuGENE transfection reagent (Promega, Madison, USA) according to the manufacturer’s instructions (1 µg/coverslip of DNA). After transfection, astrocytes were incubated at 37°C for 24 h or 48 h. In some experiments, on transfection, astrocytes were supplied with growth medium containing DGAT1 and DGAT2 inhibitors T863 (10 µM) and PF-06424439 (10 µM), respectively, to prevent biogenesis of LD, or with growth medium containing ketamine (10 µM; Tocris Bioscience, Bristol, UK) and incubated at 37°C for 24 h.

#### Immunocytochemistry and labelling of LD

After 24 or 48 h, control (non-transfected) astrocytes and astrocytes transfected with pGBW-m4134157 plasmid, encoding nsp6 protein with HA tag, were fixed with 4% formaldehyde, permeabilized with 0.1% Triton X-100 and incubated for 1 h at 37°C in 10% goat serum before being labelled with primary mouse polyclonal antibodies against HA-Tag (1:200; antibodies.com, Cambridge, UK). After overnight incubation at +4°C, coverslips were incubated for 45 min at 37°C with Alexa Fluor 488-conjugated secondary antibodies (1:600; Invitrogen, Life Technologies, Eugene, OR, USA). After washing, astrocytes were incubated for 30 min in a fluorescent marker for neutral lipids (HCS LipidTox Red, 1:500; Thermo Fisher Scientific).

Controls and astrocytes transfected with pSBL_DK186 plasmid encoding viral ORF3a fused to EGFP (3a-EGFP) were fixed with 4% formaldehyde, washed and stained with HCS LipidTOX Red. Coverslips were then mounted onto glass slides using SlowFade antifade reagent with DAPI. Every set of experiments was performed at least in duplicate.

#### Imaging, image analysis and statistical analysis

Samples were imaged with an inverted confocal microscope (LSM 800; Zeiss) with a 63×/1.4 Oil differential-interference contrast (DIC) immersion objective (Zeiss) using 488 nm (Alexa Fluor 488 and EGFP), 561 nm (HCS LipidTox Red) and 405 nm (DAPI) diode laser excitation. Emission spectra were acquired with 410–540 nm, 585–700 nm and 410–470 nm band-pass emission filters. LD content in control (non-transfected) astrocytes and astrocytes transfected with plasmids encoding for viral proteins (nsp6 or 3a) were determined based on HCS LipidTOX Red-labelled pixels after applying the 20% (nsp6) or 30% (3a) threshold of the maximum fluorescent intensity in cross-sectional areas of an individual cell, using Fiji software (LOCI; University of Wisconsin, USA). The values obtained for LD content were normalized to the average value of control (non-transfected and untreated) samples in individual experiments. The number and the perimeter of the LDs were determined using Fiji and the Analyze Particles function after applying the 20% (nsp6) or 30% (3a) threshold and signal intensity (watershed) segmentation on individual images. Then, assuming that all LDs are spherical, the LD diameter was estimated from the perimeter values using the equation *d*=*C*/*π*, where *d* is the diameter and *C* is LD perimeter. Cell area and cell perimeter were determined using Fiji software and the Measure function.

The co-localization of green (Alexa Fluor 488-immunolabelled viral protein nsp6 (nsp6) or EGFP-tagged viral protein ORF3a (3a-EGFP)) and red (HCS LipidTOX Red; LD) fluorescence pixels on individual images was determined using custom-made MATLAB software as described previously^63^. Briefly, the programme counted all red, all green, and all co-localized pixels in the image. The co-localization was calculated as a percentage of co-localized pixels compared with all red pixels. The threshold intensity for the co-localized pixels was set at 10% (26 AU) of the maximal fluorescence intensity (256 AU).

Data are presented as means±standard error of the mean. Statistical analysis was determined by ANOVA on ranks or ANOVA followed by Dunn’s test or the Holm-Sidak test; *P*≤0.05 was considered significant.

#### Fluorescence microscopy

Astrocytes were imaged by SIM (ELYRA PS.1, Zeiss), using an oil-immersion objective (Plan-Apochromat 63×/1.4 Oil DIC M27; Zeiss) to determine subcellular density and localization of SARS-CoV-2 proteins and ACE2. Whole cells were imaged by obtaining 400-nm-thick z-stacks (10–20 stacks per cell). Fluorescence was excited with 405, 488 and 561 nm laser beams and was collected through band-pass emission filters (420-480 nm, 495-575 nm and 570-650 nm, respectively) and detected with an EMCCD camera (iXon 885, Andor, Oxford Instruments, Abingdon, UK).

In the experiments where we determined the percentage of SARS-CoV-2-infected cells and cell viability imaging was performed with a confocal microscope (LSM 800; Zeiss, Oberkochen, Germany). Cells were monitored using a dry objective (20×/0.8 M27, Zeiss). Alexa Fluor 546 dye was excited with a 561-nm diode laser, and the emission was measured at 560-700 nm. DAPI was excited with a 405-nm diode laser, and emission at 400-530 nm was detected. Alexa Fluor 488 was excited with a 488-nm laser beam.

#### Clinical study

A retrospective clinical trial was conducted by including 272 patients with critical COVID-19 who received ketamine and were treated in the Intensive Care Unit (ICU) of the Department of Infectious Diseases, University Medical Center Ljubljana, Slovenia, between 12.3.2020 and 28.8.2021. During this period, circulation of the wild-type (B.1.258.17) and alpha SARS-CoV-2 variants were present^60^. The present study aimed to find out possible differences in the course of COVID-19 in critically ill patients regarding the sedation that the patients received and/or the SARS-CoV-2 variant. Patients with the respective variants were divided into two groups: those who received continuous ketamine sedation (up to 200 mg/hour) and those who received ketamine for endotracheal intubation only (dosage 1–2 mg/kg of body weight). We reviewed the patient’s demographic, clinical and virological data from medical records retrospectively. The study was approved by the National Medical Ethics Committee of the Republic of Slovenia on 20 December 2022 (#0120-449/2022/3).

### Image analysis

#### Subcellular protein densities

Image analysis was conducted using Fiji/ImageJ^63^. All cells were outlined manually based on transmission or DIC images and stored as ROIs. Fluorescence signals corresponding to immunolabelled ACE-2 were automatically thresholded using the Otsu algorithm. The threshold for images of immunolabelled viral antigens (RDRP and ENV) was set to theaverage intensity + 2 standard deviations for each cell individually. The respective signal volume was then determined using the 3D objects counter in ImageJ. Signal density was expressed as the percentage of the cell volume, with the cell volume approximated as cell area multiplied by the cell height (400 nm × the number of optical slices in a z-stack). The percentage ACE2 signal at the cell periphery was determined in the basal optical slice. Each cell was divided into central and peripheral ROIs. The peripheral ROI was 5 μm wide in an average-sized cell. Then, for each cell, the respective width of the peripheral ROI was scaled using the factor (cell area/average area)^1/2^ to account for different cell areas. The morphology of the cell nuclei was evaluated using S.F., which was determined as:

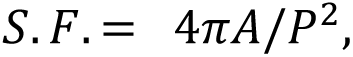

where *A* is the area and *P* is the perimeter. The morphology of cell nuclei was determined on DAPI-labelled images, which were manually thresholded, and morphometrics were obtained using the Analyze Particles function in Fiji/ImageJ.

followed by quantification of the morphology using the shape factor (S.F. = 4*πA/P*^2^; *A* indicates cell area, whereas *P* is the cell perimeter).

#### Infection rates

Cells were manually counted on DIC images overlayed with a DAPI signal. The number of cells per field of view was used as an indication of cell viability. RDRP^+^ cells were counted manually and the infection rate was expressed as the percentage of RDRP^+^ cells (of all cells) on each field of view for all experimental conditions.

## Statistical analysis

The results are presented as the means±standard error of the mean. Statistical analysis was performed in SigmaPlot 11.0 (SYSTAT, San Jose, CA, USA), GraphPad Prism (GraphPad Software, Inc., USA) and ANCOVA using MATLAB (MathWorks, Natick, MA, USA). Statistical significance was also determined with Student’s *t* test or ANOVA with the Holm-Sidak post hoc test for normally distributed data and the Mann-Whitney test or one-way ANOVA on ranks with the Kruskal-Wallis post hoc test, followed by Dunn’s method for non-normally distributed data. The fluorescence co-localization of immunolabelled proteins or membrane fluorescent probe MBF568 with 3a-EGFP and parameters of vesicle mobility (TL, MD, DI and speed) are reported as means±standard error. We also used statistical significance analysis in experiments studying LDs with the Mann-Whitney *U* test or ANOVA on ranks using SigmaPlot 11.0 (Systat Software Inc., San Jose, CA, USA), and with ANCOVA using MATLAB. Statistical significance was considered as follows: \**P*<0.05, \*\**P*<0.01 and \*\*\**P*<0.001.

## Author contributions

B.F., M.P., M.K., K.R.R., P.P., T.M.Z., J.V., S.P., P.T.V., T.S., A.H.: Investigation, Data

Analysis, Visualization, Writing – Review & Editing; K.R., N.G.K.: Clinical Data Curation, Ethical Committee Application; T.A.-Ž.: Supervision, Visualization, Writing – Review & Editing; N.V.: Conceptualization, Investigation, Data Analysis, Visualization, Supervision, Writing – Review & Editing. M.S.: Conceptualization, Investigation, Visualization, Supervision, Writing – Review & Editing; M.J.: Conceptualization, Visualization. Supervision, Writing – Review & Editing; J.J.: Conceptualization, Investigation, Visualization, Supervision, Writing – Review & Editing; R. Zorec: Directing and Overlooking the Study, Conceptualization, Investigation, Visualization, Project Administration, Supervision, Writing – Review & Editing.

## Competing interests

The authors declare no competing interests.

## Code availability

Materials and code are available from the authors on reasonable request.

## Acknowledgements

This work was supported by grants from the Slovenian Research and Innovation Agency (P3-0310, J3-2523, J3-50104, MR+, I0-0034, I0-0022), MRIC-Carl Zeiss Reference Centre for Laser Confocal Microscopy (P3-0083), MRIC UL (ICBSL3+; grant I0-0510), COST action (CA18133; ERNEST) and the European Union’s Horizon 2020 research and innovation program EVA GLOBAL project (grant agreement no. 871029). The funders had no role in the design of the study, the collection, analysis or interpretation of data, writing the manuscript or the decision to publish the results. We thank Mr. P. Runovc and Mr. M. Pate for their support in the laboratory.

## Notes

### Competing Interest Statement

The authors have declared no competing interest.

## References

1 Al-Sarraj, S. et al. Invited Review: The spectrum of neuropathology in COVID-19. Neuropathol Appl Neurobiol 47, 3–16, doi:10.1111/nan.12667 (2021).

2 Bauer, L. et al. The neuroinvasiveness, neurotropism, and neurovirulence of SARS-CoV-2. Trends Neurosci 45, 358–368, doi:10.1016/j.tins.2022.02.006 (2022).

3 Thakur, K. T. et al. COVID-19 neuropathology at Columbia University Irving Medical Center/New York Presbyterian Hospital. Brain 144, 2696–2708, doi:10.1093/brain/awab148 (2021).

4 Meinhardt, J. et al. Olfactory transmucosal SARS-CoV-2 invasion as a port of central nervous system entry in individuals with COVID-19. Nat Neurosci 24, 168–175, doi:10.1038/s41593-020-00758-5 (2021).

5 Matschke, J. et al. Neuropathology of patients with COVID-19 in Germany: a post-mortem case series. Lancet Neurol 19, 919–929, doi:10.1016/S1474-4422(20)30308-2 (2020).

6 Crunfli, F. et al. Morphological, cellular, and molecular basis of brain infection in COVID-19 patients. Proc Natl Acad Sci U S A 119, e2200960119, doi:10.1073/pnas.2200960119 (2022).

7 Verkhratsky, A. & Nedergaard, M. Physiology of Astroglia. Physiol Rev 98, 239–389, doi:10.1152/physrev.00042.2016 (2018).

8 Vander Heiden, M. G., Cantley, L. C. & Thompson, C. B. Understanding the Warburg effect: the metabolic requirements of cell proliferation. Science 324, 1029–1033, doi:324/5930/1029 [pii] 10.1126/science.1160809 (2009).

9 Escartin, C. et al. Reactive astrocyte nomenclature, definitions, and future directions. Nat Neurosci 24, 312–325, doi:10.1038/s41593-020-00783-4 (2021).

10 Verkhratsky, A. et al. Astrocytes in human central nervous system diseases: a frontier for new therapies. Signal Transduct Target Ther 8, 396, doi:10.1038/s41392-023-01628-9 (2023).

11 Domino, E. F. Taming the ketamine tiger. 1965. Anesthesiology 113, 678-684, doi:10.1097/ALN.0b013e3181ed09a2 (2010).

12 Garner, O. et al. Impact of ketamine as an adjunct sedative in acute respiratory distress syndrome due to COVID-19 Pneumonia. Respir Med 189, 106667, doi:10.1016/j.rmed.2021.106667 (2021).

13 Smischney, N. J., Beach, M. L., Loftus, R. W., Dodds, T. M. & Koff, M. D. Ketamine/propofol admixture (ketofol) is associated with improved hemodynamics as an induction agent: a randomized, controlled trial. J Trauma Acute Care Surg 73, 94–101, doi:10.1097/TA.0b013e318250cdb8 (2012).

14 Loix, S., De Kock, M. & Henin, P. The anti-inflammatory effects of ketamine: state of the art. Acta Anaesthesiol Belg 62, 47–58 (2011).

15 Irani, A. H., Steyn-Ross, D. A., Steyn-Ross, M. L., Voss, L. & Sleigh, J. The molecular dynamics of possible inhibitors for SARS-CoV-2. J Biomol Struct Dyn 40, 10023–10032, doi:10.1080/07391102.2021.1942215 (2022).

16 Nealon, J. & Cowling, B. J. Omicron severity: milder but not mild. Lancet 399, 412–413, doi:10.1016/S0140-6736(22)00056-3 (2022).

17 Islam, S., Islam, T. & Islam, M. R. New Coronavirus Variants are Creating More Challenges to Global Healthcare System: A Brief Report on the Current Knowledge. Clin Pathol 15, 2632010X221075584, doi:10.1177/2632010X221075584 (2022).

18 Knaus, W. A., Draper, E. A., Wagner, D. P. & Zimmerman, J. E. APACHE II: a severity of disease classification system. Crit Care Med 13, 818–829 (1985).

19 Potokar, M., Zorec, R. & Jorgacevski, J. Astrocytes Are a Key Target for Neurotropic Viral Infection. Cells 12, doi:10.3390/cells12182307 (2023).

20 Hernandez, V. S. et al. ACE2 expression in rat brain: Implications for COVID-19 associated neurological manifestations. Exp Neurol 345, 113837, doi:10.1016/j.expneurol.2021.113837 (2021).

21 Lasič, E. et al. Subanesthetic doses of ketamine stabilize the fusion pore in a narrow flickering state in astrocytes. J Neurochem 138, 909–917, doi:10.1111/jnc.13715 (2016).

22 Lasic, E. et al. Astrocyte Specific Remodeling of Plasmalemmal Cholesterol Composition by Ketamine Indicates a New Mechanism of Antidepressant Action. Sci Rep 9, 10957, doi:10.1038/s41598-019-47459-z (2019).

23 Stenovec, M. et al. Ketamine Inhibits ATP-Evoked Exocytotic Release of Brain-Derived Neurotrophic Factor from Vesicles in Cultured Rat Astrocytes. Mol. Neurobiol. 53, 6882–6896, doi:10.1007/s12035-015-9562-y (2016).

24 Lim, S., Zhang, M. & Chang, T. L. ACE2-Independent Alternative Receptors for SARS-CoV-2. Viruses 14, doi:10.3390/v14112535 (2022).

25 Cortese, M. et al. Integrative Imaging Reveals SARS-CoV-2-Induced Reshaping of Subcellular Morphologies. Cell Host Microbe 28, 853–866 e855, doi:10.1016/j.chom.2020.11.003 (2020).

26 Vardjan, N., Kreft, M. & Zorec, R. Dynamics of β-adrenergic/cAMP signaling and morphological changes in cultured astrocytes. Glia 62, 566–579, doi:10.1002/glia.22626 (2014).

27 Zhou, S. et al. SARS-CoV-2 E protein: Pathogenesis and potential therapeutic development. Biomed Pharmacother 159, 114242, doi:10.1016/j.biopha.2023.114242 (2023).

28 Kern, D. M. et al. Cryo-EM structure of SARS-CoV-2 ORF3a in lipid nanodiscs. Nat Struct Mol Biol 28, 573–582, doi:10.1038/s41594-021-00619-0 [pii] (2021).

29 Shariq, M., Malik, A. A., Sheikh, J. A., Hasnain, S. E. & Ehtesham, N. Z. Regulation of autophagy by SARS-CoV-2: The multifunctional contributions of ORF3a. J Med Virol 95, e28959, doi:10.1002/jmv.28959 (2023).

30 Chen, D. et al. ORF3a of SARS-CoV-2 promotes lysosomal exocytosis-mediated viral egress. Dev Cell 56, 3250–3263 e3255, doi:10.1016/j.devcel.2021.10.006 (2021).

31 Ricciardi, S. et al. The role of NSP6 in the biogenesis of the SARS-CoV-2 replication organelle. Nature 606, 761–768, doi:10.1038/s41586-022-04835-6 (2022).

32 Bills, C., Xie, X. & Shi, P. Y. The multiple roles of nsp6 in the molecular pathogenesis of SARS-CoV-2. Antiviral Res 213, 105590, doi:10.1016/j.antiviral.2023.105590 (2023).

33 Sun, X. et al. SARS-CoV-2 non-structural protein 6 triggers NLRP3-dependent pyroptosis by targeting ATP6AP1. Cell Death Differ 29, 1240–1254, doi:10.1038/s41418-021-00916-7 (2022).

34 Sun, X. et al. SARS-CoV-2 targets the lysosome to mediate airway inflammatory cell death. Autophagy 18, 2246–2248, doi:10.1080/15548627.2021.2021496 (2022).

35 Potokar, M. et al. Intermediate filaments attenuate stimulation-dependent mobility of endosomes/lysosomes in astrocytes. Glia 58, 1208–1219, doi:10.1002/glia.21000 (2010).

36 Ren, Y. et al. The ORF3a protein of SARS-CoV-2 induces apoptosis in cells. Cell Mol Immunol 17, 881–883, doi:10.1038/s41423-020-0485-9

37 Smolič, T. et al. Astrocytes in stress accumulate lipid droplets. Glia 69, 1540–1562, doi:10.1002/glia.23978 (2021).

38 Sachdev, V. et al. Novel role of a triglyceride-synthesizing enzyme: DGAT1 at the crossroad between triglyceride and cholesterol metabolism. Biochim Biophys Acta 1861, 1132–1141, doi:10.1016/j.bbalip.2016.06.014 (2016).

39 Naik, R. et al. Therapeutic strategies for metabolic diseases: Small-molecule diacylglycerol acyltransferase (DGAT) inhibitors. ChemMedChem 9, 2410–2424, doi:10.1002/cmdc.201402069 (2014).

40 Qu, Y., Wang, W., Xiao, M. Z. X., Zheng, Y. & Liang, Q. The interplay between lipid droplets and virus infection. J. Med. Virol. 95, e28967, 10.1002/jmv.28967 (2023).

41 Potokar, M., Jorgačevski, J. & Zorec, R. Astrocytes in Flavivirus Infections. Int J Mol Sci 20, doi:10.3390/ijms20030691 (2019).

42 Chauhan, A., Mehla, R., Vijayakumar, T. S. & Handy, I. Endocytosis-mediated HIV-1 entry and its significance in the elusive behavior of the virus in astrocytes. Virology 456–457, 1-19, doi:10.1016/j.virol.2014.03.002 (2014).

43 Zorec, R., Zupanc, T. A. & Verkhratsky, A. Astrogliopathology in the infectious insults of the brain. Neurosci Lett 689, 56–62, doi:10.1016/j.neulet.2018.08.003 (2019).

44 Tavcar, P. et al. Neurotropic Viruses, Astrocytes, and COVID-19. Front Cell Neurosci 15, 662578, doi:10.3389/fncel.2021.662578 (2021).

45 Mohan, J. & Wollert, T. Membrane remodeling by SARS-CoV-2 - double-enveloped viral replication. Fac Rev 10, 17, doi:10.12703/r/10-17 (2021).

46 Gioia, U. et al. SARS-CoV-2 infection induces DNA damage, through CHK1 degradation and impaired 53BP1 recruitment, and cellular senescence. Nat Cell Biol 25, 550–564, doi:10.1038/s41556-023-01096-x (2023).

47 Ghosh, S. et al. beta-Coronaviruses Use Lysosomes for Egress Instead of the Biosynthetic Secretory Pathway. Cell 183, 1520–1535 e1514, doi:10.1016/j.cell.2020.10.039 (2020).

48 Wang, W. et al. Genetic variety of ORF3a shapes SARS-CoV-2 fitness through modulation of lipid droplet. J. Med. Virol. 95, e28630, 10.1002/jmv.28630 (2023).

49 Dong, H. & Czaja, M. J. Regulation of lipid droplets by autophagy. Trends Endocrinol. Metab. 22, 234–240, 10.1016/j.tem.2011.02.003 (2011).

50 Yuan, Z. et al. ATG14 targets lipid droplets and acts as an autophagic receptor for syntaxin18-regulated lipid droplet turnover. Nature Communications 15, 631, doi:10.1038/s41467-024-44978-w (2024).

51 Nardacci, R. et al. Evidences for lipid involvement in SARS-CoV-2 cytopathogenesis. Cell Death Dis. 12, 263, doi:10.1038/s41419-021-03527-9 (2021).

52 Dias, S. S. G. et al. Lipid droplets fuel SARS-CoV-2 replication and production of inflammatory mediators. PLoS Path. 16, e1009127, doi:10.1371/journal.ppat.1009127 (2020).

53 Schwartz, J. P. & Wilson, D. J. Preparation and characterization of type 1 astrocytes cultured from adult rat cortex, cerebellum, and striatum. Glia 5, 75–80, doi:10.1002/glia.440050111 (1992).

54 Quintero, M., Cabanas, M. E. & Arus, C. A possible cellular explanation for the NMR-visible mobile lipid (ML) changes in cultured C6 glioma cells with growth. Biochim Biophys Acta 1771, 31–44, doi:10.1016/j.bbalip.2006.10.003 (2007).

55 Rigal, A., Doyle, S. M. & Robert, S. Live cell imaging of FM4-64, a tool for tracing the endocytic pathways in Arabidopsis root cells. Methods Mol Biol 1242, 93–103, doi:10.1007/978-1-4939-1902-4_9 (2015).

56 Pirnat, S. et al. Astrocyte arborization enhances Ca(2+) but not cAMP signaling plasticity. Glia 69, 2899–2916, doi:10.1002/glia.24076 (2021).

57 Grynkiewicz, G., Poenie, M. & Tsien, R. Y. A new generation of Ca2+ indicators with greatly improved fluorescence properties. J Biol Chem 260, 3440–3450 (1985).

58 Potokar, M., Kreft, M., Pangrsic, T. & Zorec, R. Vesicle mobility studied in cultured astrocytes. Biochem Biophys Res Commun 329, 678–683, doi:S0006-291X(05)00283-4 [pii]10.1016/j.bbrc.2005.02.030 (2005).

59 Stenovec, M., Bozic, M., Pirnat, S. & Zorec, R. Astroglial Mechanisms of Ketamine Action Include Reduced Mobility of Kir4.1-Carrying Vesicles. Neurochem Res, doi:10.1007/s11064-019-02744-1 (2019).

60 Suljic, A. et al. Efficient SARS-CoV-2 variant detection and monitoring with Spike Screen next-generation sequencing. Brief Bioinform 25, doi:10.1093/bib/bbae263 (2024).

61 Lei, C., Yang, J., Hu, J. & Sun, X. On the Calculation of TCID(50) for Quantitation of Virus Infectivity. Virol Sin 36, 141–144, doi:10.1007/s12250-020-00230-5 (2021).

62 Kreft, M., Milisav, I., Potokar, M. & Zorec, R. Automated high through-put colocalization analysis of multichannel confocal images. Computer methods and programs in biomedicine 74, 63–67, doi:10.1016/s0169-2607(03)00071-3 (2004).

63 Schindelin, J. et al. Fiji: an open-source platform for biological-image analysis. Nat Methods 9, 676–682, doi:10.1038/nmeth.2019 (2012).

