## Supplementary data for "Ketamine inhibition of SARS-CoV-2 replication in astrocytes is associated with COVID-19 disease severity in a variant-dependent manner"

### SUPPLEMENTARY FIGURES

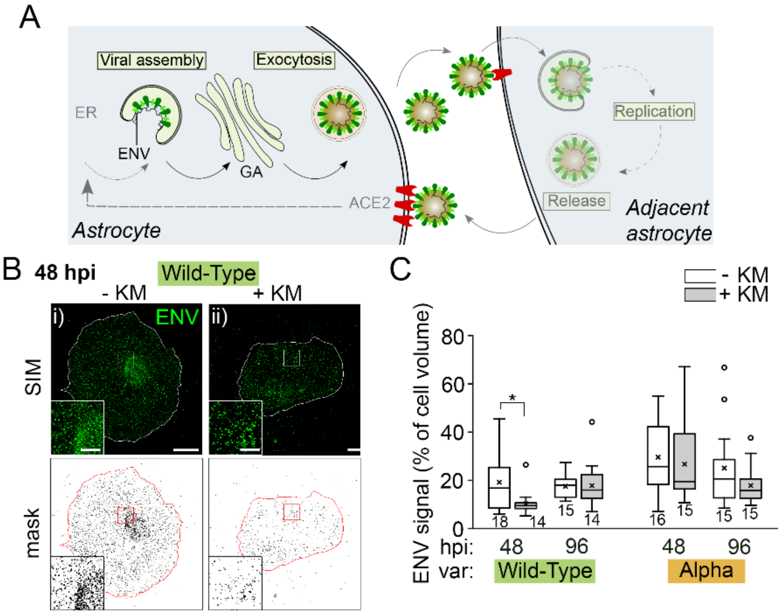


**Supplementary Fig. 1. Ketamine treatment of wild-type (B.1.258.17) infected human astrocytes decreases the subcellular density of the envelope viral (ENV) protein**. (**A**) The schematic illustrates the mechanism of viral release from human astrocytes, mediating the infection of adjacent cells. (**B**) Fluorescent micrographs recorded by structured illumination microscopy (SIM) show astrocytes immunolabelled against the envelope protein (ENV) of SARS-CoV-2 by anti-ENV antibody (see Methods). "Masks" below the micrographs are shown in black (signal) and white (background). Similarly, for the RDRP signal (Fig. 5), we observed a reduction in the ENV signal at 48 hpi in ketamine-treated astrocytes infected with the wild-type variant ($\mathbf{B_i}$ versus $\mathbf{B_{ii}}$). (**C**) This reduction was confirmed with quantitative analysis of the viral protein signal expressed as the percentage relative to the cell volume ($P$=0.035, Mann-Whitney $U$ test). The numbers below the boxplots are the number of cells. GA, Golgi apparatus; hpi, hours post-infection; KM, ketamine; var, variant. Data are from two human donors. Scale bar, 20 µm (insets, 5 µm).

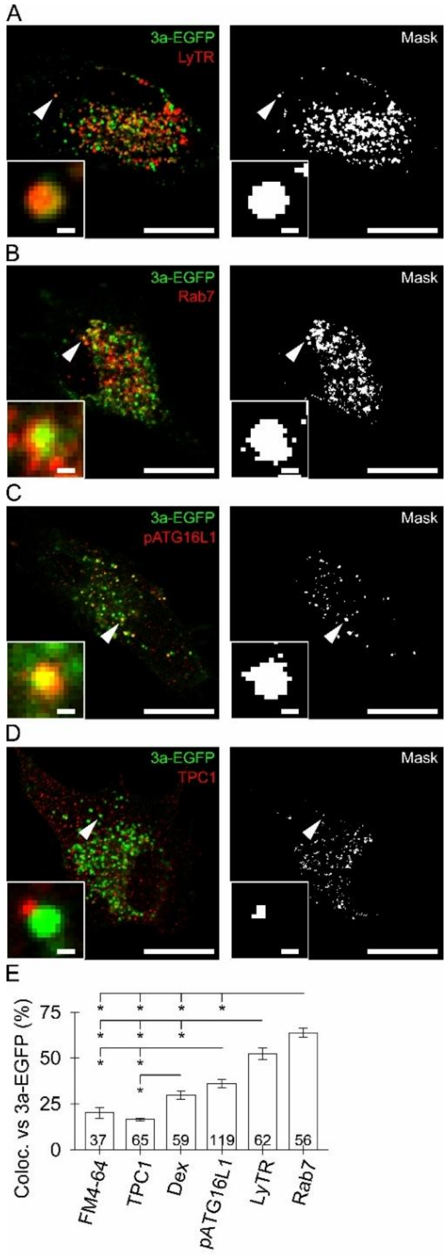


**Supplementary Fig. 2. Protein 3a-EGFP predominantly segregates to acidified late endolysosomes and to early autophagosomes**. (**A–D**) Confocal images of 3a-EGFP-positive vesicles (green fluorescent puncta) in transfected astrocytes stained with LysoTracker Red DND-99 (LyTR), which labels acidified vesicles (red, **A**), with immunolabelled Rab7, a marker of late endolysosomes (red, **B**), with immunolabelled pATG16L1, a marker of early autophagosomes (red, **C**), and with immunolabelled TPC1, characteristic of early and recycling endosomes (red, **D**). The mask images (white) display co-localized pixels (**A–D**, right) and insets display magnified views of the selected vesicles (arrowheads). Scale bars, 20 μm and 0.5 μm (insets) (**A–D**). (**E**) Quantitative co-localization (mean±standard error; %) of fluorescent FM4-64, TPC1, pATG16L1, LyTR and Rab7 with 3a-EGFP fluorescence. Note the substantial co-localization between LyTR and 3a-EGFP and between Rab7 and 3a-EGFP, indicating that 3a-EGFP predominantly segregated into acidified late endolysosomes. The numbers at the bottom of the bars are the number of cell images analysed. *$P<0.05$ versus respective comparison (ANOVA on ranks followed by Dunn's post hoc test).

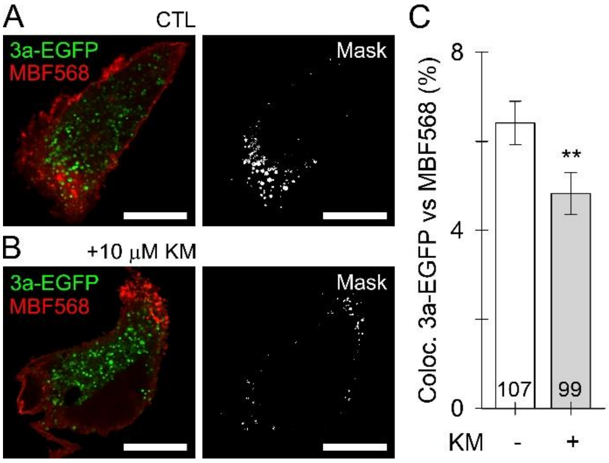


**Supplementary Fig. 3. Ketamine treatment reduces the translocation of the protein 3a-EGFP to the astrocyte surface. (A, B)** Confocal images of 3a-EGFP vesicles in non-treated control (CTL) and ketamine-treated (+10 µM KM) astrocytes labelled with MemBrite Fix 568/580 (MBF568 red, **A**, **B**), a marker of the plasmalemma. The mask images (white) display co-localized pixels (**A**, **B**, right). Scale bars, 20 µm (**A**, **B**). **(C)** Quantitative fluorescence co-localization (mean±standard error; %) of 3a-EGFP with the membrane dye MBF568 in non-treated controls (CTL) and ketamine-treated astrocytes (+10 µM KM), respectively. Note ketamine-mediated reduced protein 3a-EGFP localization at the astrocyte surface. The numbers at the bottom of the bars are the number of cell images analysed (**C**); $^{**}P<0.01$ versus control (Mann-Whitney *U* test).

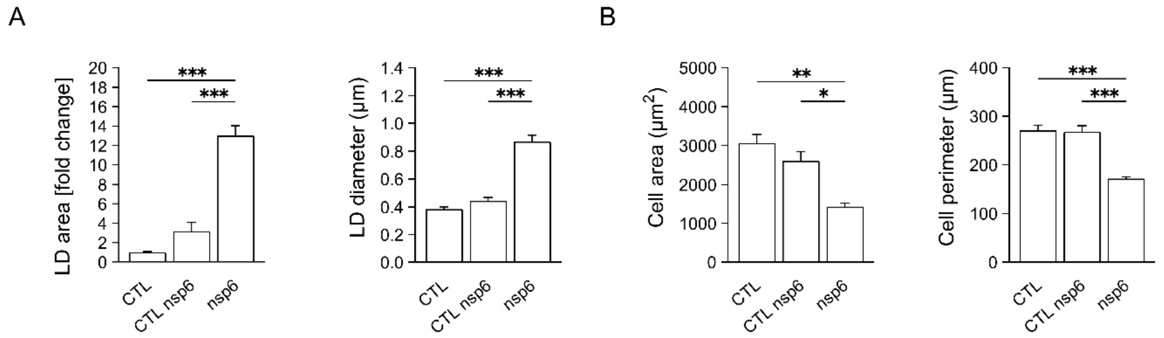


**Supplementary Fig. 4. Prolonged (48 h) overexpression of viral nsp6 protein in rat cortical astrocytes increases the accumulation of lipid droplets and reduces cell size**. (**A**) Histograms of mean LipidTOX-stained cell area normalized to control untreated samples (lipid droplet [LD] area [fold change]), mean diameter of LipidTOX-stained LDs (LD diameter), and (**B**) mean cell area and mean cell perimeter in cortical astrocytes in non-transfected cell cultures (CTL; *n*=3; 39 cells) and in cell cultures transfected with pDNA encoding the viral nsp6 protein, in cells without apparent nsp6 expression (CTL nsp6; *n*=5; 64 cells) and in cells expressing nsp6 (nsp6; *n*=7; 76 cells). To trace nsp6 expression and LD formation, cultures were immunocytochemically labelled with antibodies against nsp6 and the fluorescent LD marker HCS LipidTOX Red (red) 48 h on transfection. Data are presented as means±standard error of the mean. *n*, number of independent experiments. *$P<0.05$, **$P<0.01$, ***$P<0.001$ all pairwise (ANOVA, Holm-Sidak test [LD area, LD diameter, cell perimeter] and ANOVA on ranks, Dunn's test [cell area]).

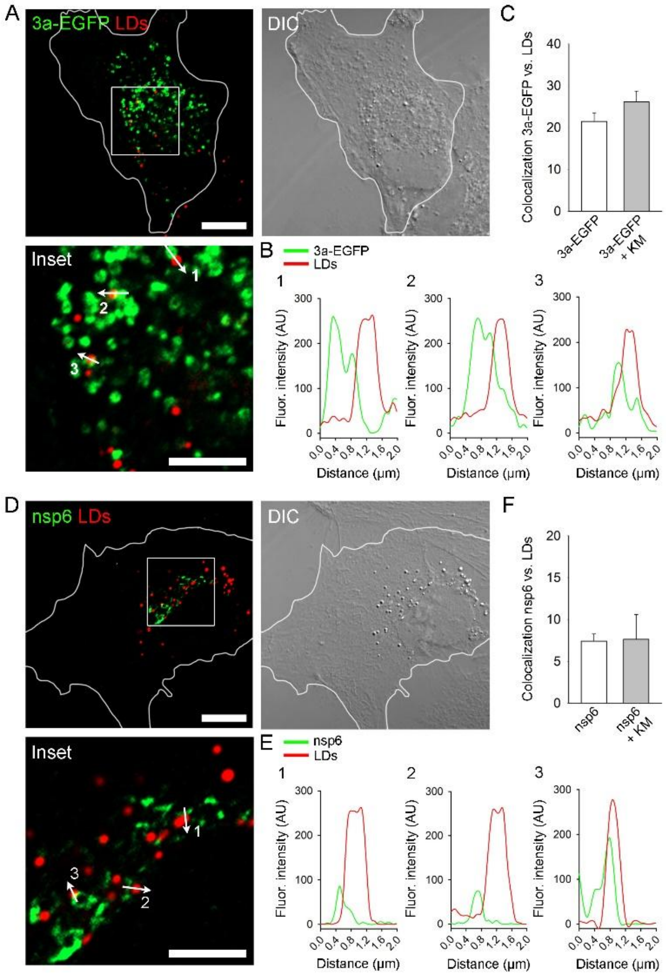


**Supplementary Fig. 5. Ketamine does not affect the level of co-localization of viral proteins nsp6 and 3a with lipid droplets in cortical astrocytes**. (**A**, **D**) Representative fluorescence images of isolated cortical astrocytes transfected with pDNA encoding the viral EGFP-tagged 3a protein (**A**) and nsp6 protein (**D**). To trace nsp6 expression, cell cultures were immunocytochemically labelled first with primary antibodies against HA-Tag on nsp6 and then with anti-HA Alexa Fluor 488 secondary antibodies. Lipid droplets (LDs) were labelled with the fluorescent LD marker HCS LipidTOX Red (red) 24 h after transfection. The same images taken by differential-interference contrast microscopy (DIC) are shown in the upper right panel). The white line shows the outline of the cells. Insets (lower left panels) show white boxed regions from the upper left panels at higher magnification. Scale bars, 10 µm and 5 µm (inset). (**B**, **E**) Fluorescence intensity profiles of green fluorescent signal (3a-EGFP (**B**) and nsp6 (**E**)) and red fluorescent signal (LDs) along the arrow lines 1, 2 and 3 in the respective insets. (**C**, **F**) Mean co-localization (%) of green fluorescent pixels (3a-EGFP (**C**) and nsp6 (**F**)) versus red fluorescent pixels (LDs) at 20% (3a-EGFP) or 10% (nsp6) threshold of the maximal fluorescence intensity in untreated astrocytes expressing 3a-EGFP (*n*=7; 96 cells) or nsp6 (*n*=15; 249 cells) and in astrocytes expressing 3a-EGFP or nsp6 on 24 h exposure to 10 µM ketamine (3a-EGFP+KM; *n*=6; 60 cells or nsp6+KM; *n*=4; 53 cells).

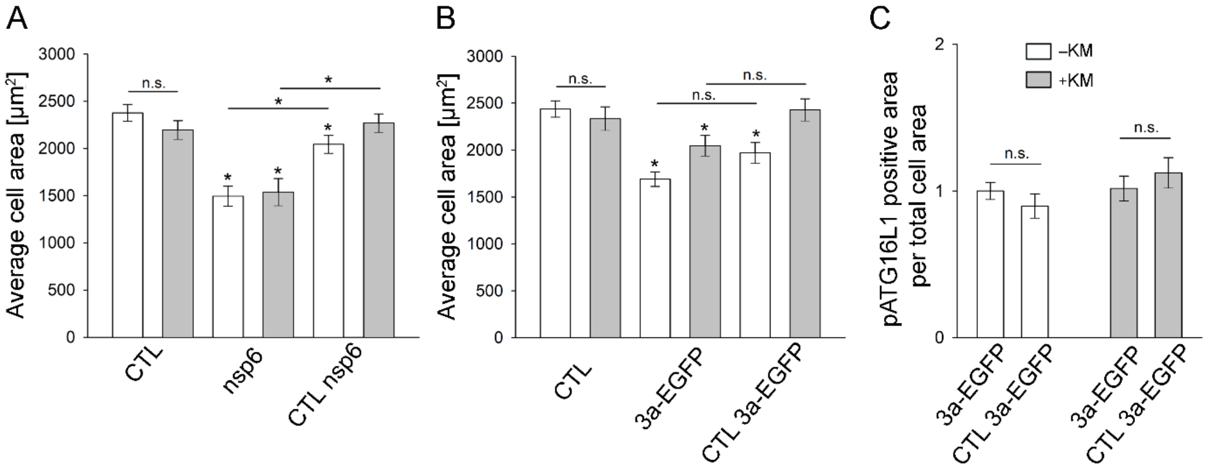


**Supplementary Fig. 6. Ketamine treatment does not affect the average cell size (cell area) in control, transfected cells by 3a-EGFP and nsp6 proteins and in non-transfected cells. (A)** Mean reduction in cell area in astrocytes transfected by nsp6 is not sensitive to ketamine treatment. **(B)** Mean reduction in cell area in astrocytes transfected by 3a is not sensitive to ketamine treatment. **(C)** Control experiments demonstrate that transfection of EGFP does not affect the mean cell area measurements.
